# Dispersal, adaptation and persistence of H5N1 in the sub-Antarctic and Antarctica

**DOI:** 10.64898/2026.03.20.713283

**Authors:** Augustin Clessin, Marius Brusselmans, Samuel L. Hong, Jérémy Tornos, Mathilde Lejeune, Yucai Shao, François-Xavier Briand, Celia Abolnik, Benjamin Kaza, Marc A. Suchard, Begoña Aguado, Antonio Alcamí, Christophe Barbraud, Martin Beer, Ashley Bennison, Francesco Bonadonna, Timothée Bonnet, Charles-André Bost, Sara Boucheron, Tristan Bralet, Paulo Catry, Jaimie Cleeland, Maelle Connan, Holly A. Coombes, Karine Delord, Camille de Pasquale, Meagan L. Dewar, Xiaomin Dong, Julia Emerit, Romain Fischer, Nicholas Fountain-Jones, Filippo Galimberti, Jacob González-Solís, Christophe Guinet, Carrie Gunn, Anne Günther, Matteo Iervolino, Joe James, Xiang Ji, Ian Jonsen, Christopher W. Jones, Amanda Kuepfer, Thijs Kuiken, Céline Lebohec, Simeon Lisovski, Lucia Llorente Zubiri, Joshua G. Lynton-Jenkins, Paula Martinez-García, Michelle McCulley, Clive R. McMahon, Benjamin C. Mollett, Ana I. Moraga-Quintanilla, Rhiannon Nichol, Aude Noiret, Maria Ogrzewalska, Kate Owen, Catalina Pardo-Roa, Samuel Peroteau, Richard A. Phillips, Elie Poulin, Andrew Rambaut, Scott M. Reid, Eva Riehle, Michelle Risi, Odin Rumianowski, Peter G. Ryan, Simona Sanvito, Andrew Stanworth, Antje Steinfurth, Jonathon Stevenson, Antoine Stier, Marcela M. Uhart, Ralph E. T. Vanstreels, Angela Vázquez-Calvo, Juliana A. Vianna, Katie Wells, Jeff White, Paul Whitelaw, Michelle Wille, Jane Younger, Laura C. Roberts, Béatrice Grasland, Ashley C. Banyard, Martha I. Nelson, Amandine Gamble, Thierry Boulinier, Guy Baele

## Abstract

High pathogenicity avian influenza virus (HPAIV) H5N1 reached the sub-Antarctic and Antarctica in 2023, subsequently spreading to remote locations within this region where it had devastating impacts on seal, penguin and albatross populations. The threat to marine wildlife over this broad area exemplifies the need to understand H5N1 long-distance dispersal and evolution. We obtained 104 novel viral genomic sequences from samples that we collected at South Georgia, Kerguelen, Crozet, Prince Edward, Falklands/Malvinas Islands and the Antarctic Peninsula in a region spanning 8,000 kilometers. Using recent phylogeographic modeling advances we show that H5N1 spread encompassed numerous transmission events between distant locations, accumulating mammalian-adaptive mutations in the process. Seals are the most affected species, but we reveal that the long-distance eastward virus dispersal better aligns with the long-distance movements of large petrels and albatrosses. The risk of H5N1 endemisation, dispersal to other locations and ongoing evolution are highly concerning.

## Main

Due to their isolation from other continents, the sub-Antarctic and Antarctica may be considered at low risk of exposure to epizootic infectious agents ^1^. Prior to September 2023, high pathogenicity avian influenza viruses (HPAIVs) had never been detected in the sub-Antarctic and Antarctica, despite several epizootic waves that spread across most continents during recent decades ^2,3^. However, this isolation is not absolute as low pathogenicity avian influenza viruses have previously been documented in the Antarctic Peninsula and their close genetic proximity with South American strains clearly illustrates the existence of some connectivity ^4–7^.

Following the spread of HPAIV H5N1 clade 2.3.4.4b along the coasts of South America in 2022 and 2023 ^8–12^, several seabird and pinniped species that migrate to the sub-Antarctic and Antarctica had been found infected, leading to growing concerns about the potentially devastating effects that the virus could have on wildlife if it reached these regions ^2,13^. The virus was eventually detected on South Georgia Island / Islas Georgias del Sur (hereafter South Georgia), and the Falkland Islands / Islas Malvinas (hereafter the Falklands) in the sub-Antarctic in September and October 2023, i.e., at the beginning of the austral summer when most seabird and seal species return to their breeding grounds ^2,14^. In January 2024, HPAIV H5N1 was detected on the Antarctic Peninsula ^15,16^. At the beginning of the next austral summer, in September and October 2024, the virus was detected on Gough Island (hereafter Gough) in the Atlantic Ocean, and on Marion Island in the Prince Edward archipelago (hereafter Prince Edward), Possession Island in the Crozet archipelago (hereafter Crozet) and Kerguelen Islands (hereafter Kerguelen) in the southern Indian Ocean ^17–19^. The virus was still reported in the austral summer 2024/25 at the Falklands, South Georgia and the Antarctic Peninsula, but still nowhere else in the Antarctic continent ^18,20,21^.

Given the remoteness of these locations, assessing the complete impact of HPAIV H5N1 on the populations of sub-Antarctic and Antarctic seabirds and marine mammals is challenging, and the long-term impacts remain unknown. However, current evidence points towards a dramatic impact on South Georgia, where about 47% of the breeding female southern elephant seals (*Mirounga leonina*) – some 53,000 individuals – are suspected to have died from HPAIV H5N1 during the austral summer 2023/24 ^22^. In the Falklands, black-browed albatrosses (*Thalassarche melanophris*) recorded the highest number of deaths, potentially up to a hundred thousand, mostly chicks ^23^. These numbers would make the Falklands and South Georgia the two archipelagos with the highest numbers of fatalities, by at least one and perhaps two orders of magnitude. On the Antarctic Peninsula, previous reports suggest hundreds of dead skuas (*Stercorarius* spp.) and crabeater seals (*Lobodon carcinophaga*) ^15,21,24,25^. In the Indian Ocean, the first publications suggest mortality numbers in the thousands – mainly southern elephant seals but also king penguins (*Aptenodytes patagonicus*) and snowy albatrosses (*Diomedea exulans*) ^17,26^. Notwithstanding the absence of comprehensive assessment of mortality, the available data point toward hundreds of thousands of wildlife deaths and urges the need to understand HPAIV H5N1 transmission pathways, dynamics and spread mechanisms in the sub-Antarctic and Antarctica.

Wild bird movements are important drivers of the spread of several HPAIVs ^27–29^. Among migratory birds, some seabird species perform long migrations and can spread pathogens across hemispheres, oceans and between continents ^29^. The distance and direction of their movements depend on the species, animal age, season and individual preference, all of which need to be considered when generating hypotheses about how movement and disease spread are related ^30–34^. Bearing this in mind, the spread of a virus by hosts also depends on their behaviour. Whether a host species travels with or without landing on distant islands, interacts at sea with conspecifics or other species, and scavenges on land or on floating carcasses (Supplementary Figures S1-S2) are all likely to influence virus transmission in the sub-Antarctic and Antarctic regions ^35–37^. The severity and types of clinical signs, the duration of the infectious period and the tropism of the virus are host-species dependent and condition their abilities to spread the virus ^38^.

Seabird and seal presence on land is seasonal in the sub-Antarctic and Antarctica. Most species are summer breeders and their presence on land spans from September to April. During that period, many colonial species breed and molt at high densities, in single- or multi-species colonies and molting sites. In either case, predator and scavenger species abound in the surroundings. These seasonal spatial aggregations provide optimal conditions for pathogen spread between conspecifics, and numerous opportunities for inter-species transmission events, e.g. through scavenging, environmental contamination or through close contact between individuals of different species ^2,35^. During the rest of the year, most species remain at sea – densities during these periods are lower and correspondingly the contact rates between individuals are lower. A few species are winter breeders, and some breeding cycles span an entire year so some birds remain on land during winter ^39^. Among those, king penguins and snowy albatrosses are the only species reported as being affected by HPAIV H5N1. Gentoo penguins (*Pygoscelis papua*) - although summer breeders – remain local, social, and come on land during the winter as well ^40^. Other species reported to die from HPAI in the sub-Antarctic and Antarctica are mostly absent from land in winter. We hence hypothesised that the behavioural ecology of the different host species – particularly the seasonality of their breeding biology, movements, scavenging and social behaviour – would be key determinants of the dispersal of the virus, its potential persistence in the sub-Antarctic and Antarctica and its evolution.

The pandemic risk caused by influenza A viruses is well documented ^41^, and the World Health Organisation (WHO) regularly updates a public health research agenda for influenza ^42^ to promote pandemic preparedness. Central to being adequately prepared is the ongoing research effort to identify the presence and impact of mutations on the phenotype of the virus, and particularly the zoonotic risk ^43,44^. Therefore, many mutations – particularly in the polymerase complex and in the haemagglutinin – have been shown to impact the virus’s transmission risk to humans, e.g., by increasing airborne transmission risk for mammals ^45^ or increasing the capacity to replicate in mammalian cells ^46^. The ongoing outbreak of H5N1 in US dairy cattle that started in 2024 ^47^ and the H5N1 marine mammal outbreak that started in 2023 in South America ^8^ and continues in the sub-Antarctic and Antarctica both featured new PB2 mammalian adaptations (M631L and Q591K/D701N, respectively) that helped avian viruses – all segments are of avian origin – replicate and transmit efficiently in mammals over sustained time periods. The possibility to investigate the long-term transmission and evolution of an avian-origin HPAIV H5N1 spreading between mammals is new and constitutes a critical opportunity to understand the long-term drivers of mammalian adaptation of influenza viruses. Here, we present a large-scale phylogeographic analysis to infer and interpret the circumpolar spread of HPAIV H5N1 across the sub-Antarctic and Antarctica. We include all available genomic sequences from these two regions that were previously published ^14,17,19,20,24,48–51^, together with a large set of 104 novel sequences, including 5 from Prince Edward, 42 from the Falklands, 31 from South Georgia, 14 from Crozet, 7 from Kerguelen and 5 from the Antarctic Peninsula (Figure 1). Overall, these sequences represent all locations where HPAIV H5N1 has been confirmed in the sub-Antarctic and Antarctica up to March 2025. Our study raises critical concerns around the ongoing evolution of the virus, its long-term persistence in the sub-Antarctic and Antarctica, and the risk of continued spread to other locations around Antarctica and the Southern Ocean as well as to neighbouring continents.

**Figure 1.**
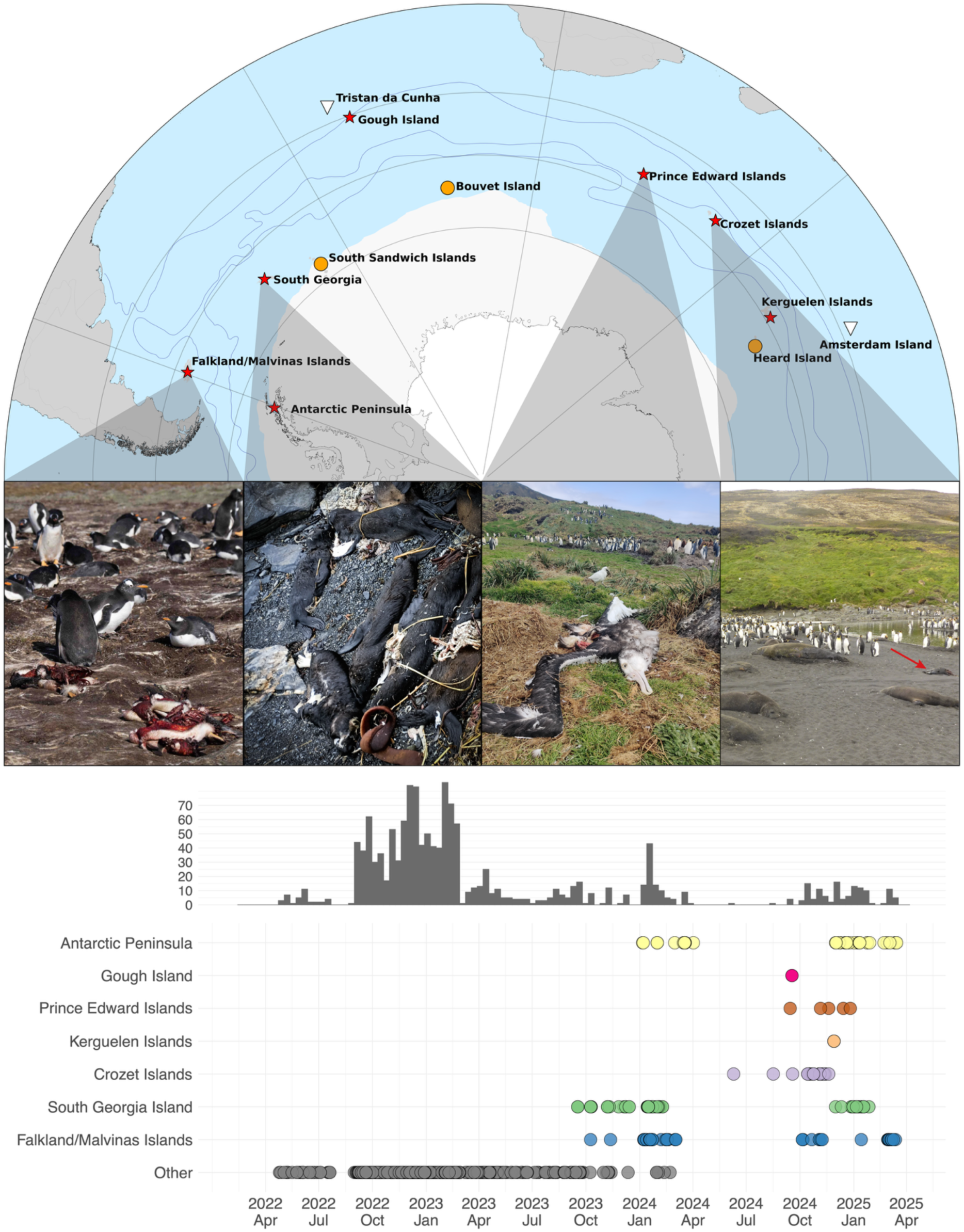
Top: map of the areas affected by HPAIV H5N1 in the Sub-Antarctic and Antarctica. Islands where HPAIV H5N1 was confirmed are represented by red stars, islands where HPAIV has not been detected are represented by a white triangle, and islands that have not been visited between September 2023 and March 2025 are represented by orange circles. Within the Antarctic Treaty Area Boundary (i.e., the Antarctic continent and adjacent islands below 60°S latitude), HPAIV H5N1 has been confirmed only in the Antarctic Peninsula and the adjacent islands. Middle: illustrations of some species affected by mass mortality events, from left to right: gentoo penguin carcasses in a colony on the Falklands (picture credit: Derek Peterson); dead Antarctic fur seal pups (*Arctocephalus gazella*) on South Georgia (picture credit: Augustin Clessin); dead juvenile snowy albatross on Prince Edward, with a live black-faced sheathbill (*Chionis minor*) and king penguins in the background (picture credit: Laura Roberts); dead elephant seals (none are alive) – with one dead king penguin (red arrow) and live king penguins in the background – on Crozet (picture credit: Jérémy Tornos & Mathilde Lejeune). Bottom: locations and dates of sampling for the HPAIV H5N1 sequences used in our dataset. Rows show the collection dates of genomes on the bottom, as well as the number of genomes as a bar plot at the top. Histogram bars use 2-week bins.

## Results

### Phylogenetic analysis uncovers Sub-Antarctic and Antarctic clades

We conducted phylogenetic and phylogeographic analyses on a dataset composed of 104 novel sequences from the sub-Antarctic and Antarctica and 1,279 background sequences, previously selected as a large monophyletic clade encompassing all sequences from the sub-Antarctic and Antarctica ^17^. All sequences belong to genotype B3.2. We first performed a time-calibrated Bayesian phylogenetic analysis, which showed the presence of two distinct clades from the sub-Antarctic and Antarctica (Figure 2, Supplementary Figures S3-S5). One clade (which we refer to as “Clade I”) descends from viruses that primarily circulated within avian hosts in South America, while the other (which we refer to as “Clade II”) descends from viruses that primarily circulated within mammalian hosts in South America and carry the Q591K and D701N mammalian adaptations in PB2. Neither Clade I nor Clade II appear to be mammalian-or avian-specific in the sub-Antarctic and Antarctica.

**Figure 2.**
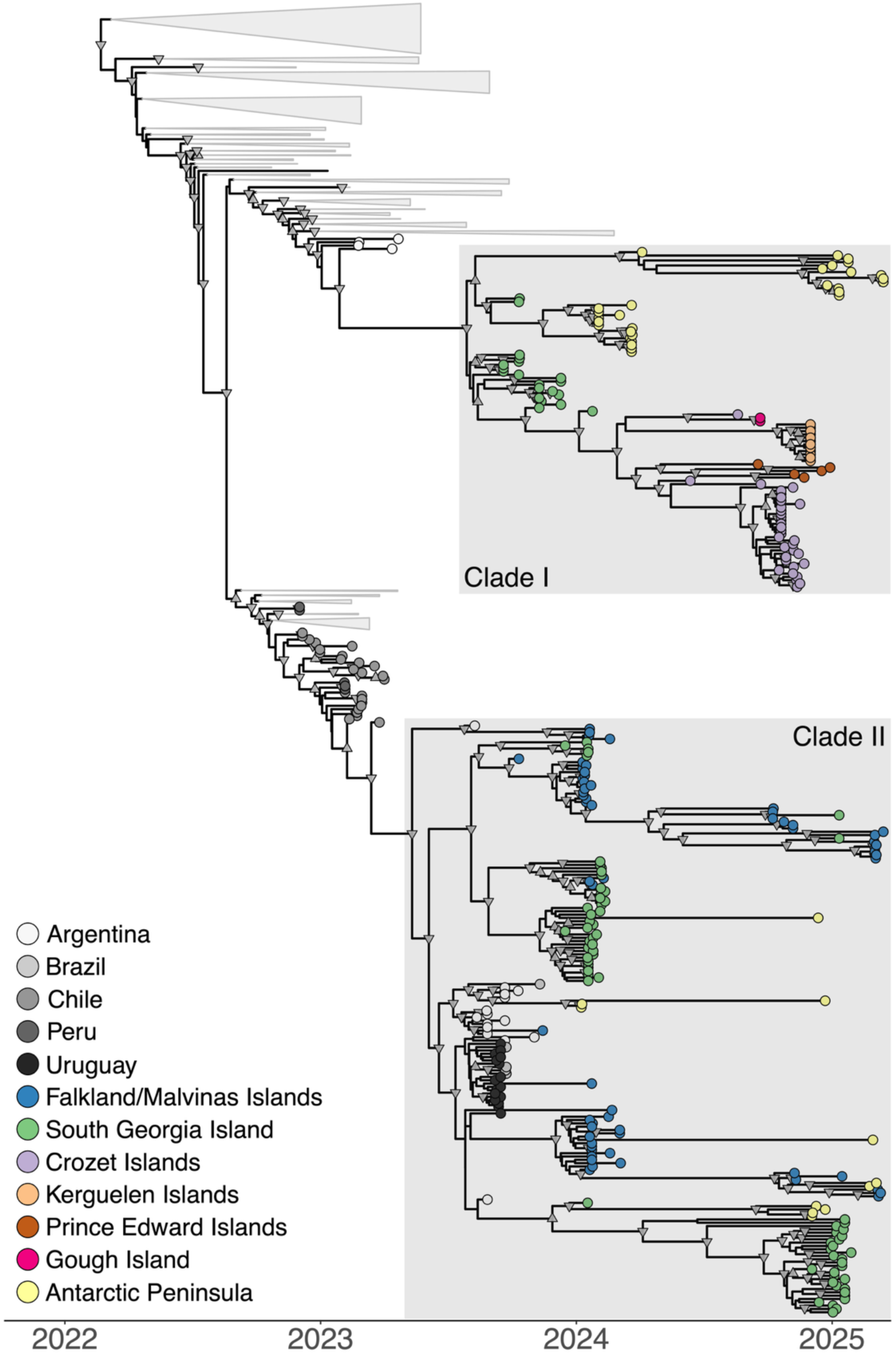
Time-calibrated consensus phylogeny of our 1383-taxa sub-Antarctic and Antarctic dataset – with novel genomic sequences from Crozet, Kerguelen, Prince Edward, South Georgia, the Falklands and the Antarctic Peninsula – identifies two sub-Antarctic and Antarctica clades. Tip colors correspond to sampling locations, and internal nodes with posterior support above 0.7 and 0.9 are marked with down-facing and up-facing triangles, respectively. Clades without any sub-Antarctic samples have been collapsed for visual clarity. Sequences from Crozet, Kerguelen, Prince Edward and Gough islands all fall within Clade I, while sequences from the Falklands cluster within Clade II.

We subsequently performed a forward-in-time (FIT) phylogeographic analysis ^52^ on an empirical tree distribution stemming from the phylogenetic analysis (Figure 2). Given the sensitivity of the FIT model to sampling bias, we also conducted a backward-in-time (BIT) phylogeographic analysis which is less sensitive to sampling bias ^53,54^. For computational efficiency, this BIT analysis was conducted independently on Clade I and Clade II. In the following paragraphs, the results are those obtained under the BIT analysis unless explicitly mentioned. Our phylogenetic results identified four possible reassortment events in South Georgia (Supplementary Figures S6 and S7), but these would have occurred between closely related lineages and only at tip nodes in our phylogeny, and we hence do not consider these to have a meaningful impact on our phylogeographic reconstruction.

### Initial introductions and spread of HPAIV H5N1 within the 2023/24 austral summer

Following the initial HPAIV detection on South Georgia in September 2023 and the Falklands in October 2023, HPAIV-related mortality of both seabirds and seals was reported continuously from South Georgia until the end of the austral summer ^55^. In contrast, on the Falklands our first confirmed report was a lone case in October 2023, and only in December 2023 were higher levels of mortality detected ^56^. HPAIV H5N1 was later confirmed in the Antarctic Peninsula in January 2024 ^13,20^. Our phylogeographic analyses (see Figure 3, Supplementary Figures S8 and S9), in line with previously published analyses, suggest that the first detected South Georgia and Falklands cases resulted from different introductions from South America, and that the first introduction to Antarctica came from South Georgia ^20^. We here estimate that the first detected cases at South Georgia and Antarctica originated in Clade I, that primarily circulated in avian hosts in South America, while the initial introduction to the Falklands originated in Clade II, that mostly circulated within marine mammals in South America ^11,12,57,58^. However, our results suggest that even though the first cases at South Georgia are within Clade I, Clade II could have been introduced earlier at South Georgia, and have remained unnoticed for several months there while being the source of the first introduction to the Falklands. The novel sequences used in our study demonstrate that, on the Falklands, at least three additional introductions of the virus occurred later that season from South America, all in Clade II. In South Georgia, a third introduction from South America was detected later in the season, also in Clade II. We estimate one additional introduction in Clade II from South America to the Antarctic Peninsula to have occurred. Following these initial introductions into the sub-Antarctic and Antarctica, further spread took place within the region. Two more transmission events are estimated to have occurred from South Georgia to the Antarctic Peninsula – one in Clade II and a second one in Clade I, and two transmission events in Clade II from South Georgia to the Falklands. In the Falklands, only seabirds were affected, mostly black-browed albatrosses. In South Georgia, various seabird species were affected throughout the season, but southern elephant seals and Antarctic fur seals were affected as well, in higher numbers than seabirds. In Antarctica, skuas were the most affected.

**Figure 3.**
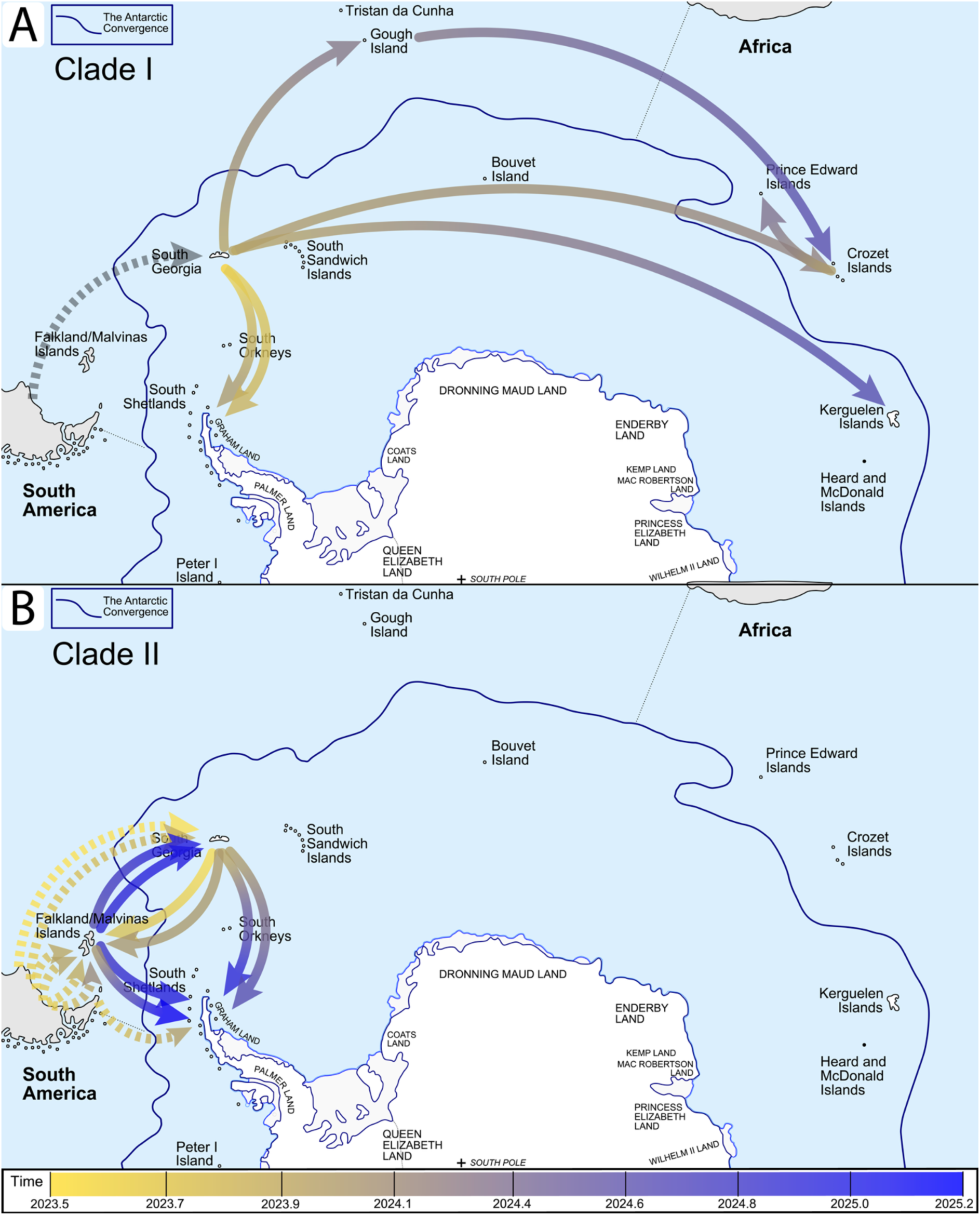
Backward-in-time phylogeographic reconstruction ^53,54^ of HPAIV H5N1 2.3.4.4b dispersal in the sub-Antarctic and Antarctica. A) After an initial introduction of Clade I from South America to South Georgia (in grey; before the start of our time line), we estimate a consistent long-distance circumpolar eastward movement in agreement with common seabird movement data (Figure 4); B) Back-and-forth dispersal of Clade II between South Georgia and the Falklands, along with introductions into Antarctica from both islands as well as from South America. The Falklands recorded multiple introductions from South America. All phylogeographic estimates are based on the currently available genomic data; comparisons with the classic FIT phylogeographic reconstruction ^52^ are shown in Supplementary Figures S11-S14. Arrows shown represent estimated single introductions and are coloured according to a time-dependent gradient. Background map adapted from ©Hogweard (talk · contribs), CC BY-SA 3.0 <https://creativecommons.org/licenses/by-sa/3.0>, via Wikimedia Commons.

### Long-distance spread of Clade I to the southern Indian Ocean

In the following breeding season (from September 2024 onward), unusual mortalities were reported further east, up to the Southern Indian Ocean – thousands of kilometers eastward – at Prince Edward and Gough (September 2024), Crozet (October 2024) and Kerguelen (November 2024). On Gough, only a few individual skuas were found dead. On Crozet and Kerguelen, mostly southern elephant seals were affected, together with some brown skuas, kelp gulls and king penguins (Crozet only). On Prince Edward, only unusual mortality among seabirds was observed, mostly of snowy albatrosses and skuas. All were confirmed as HPAIV-related. Later, we retrospectively tested and sequenced samples collected as part of the long-term mortality monitoring program implemented at Crozet. We report and provide sequences of HPAI-positive samples collected from carcasses as early as June and August 2024, i.e. five months before the beginning of the mass mortality event. Those carcasses were isolated, and did not trigger any HPAIV suspicions at the time given that the number did not exceed the mortality levels usually observed on Crozet.

Previous phylogeographic analyses initially suggested that the introductions to Crozet, Kerguelen and Gough were all independent introductions from South Georgia ^17,59^. Here, our dataset is supplemented with a large number of novel sequences from Prince Edward, South Georgia, Crozet, the Falklands and Kerguelen. Our novel analyses here confirm that the infections from Kerguelen, Crozet and Gough all belong to Clade I (Figure 2) and additionally demonstrate that the novel sequences from Prince Edward also belong to that clade. However, the improved resolution of phylogenetic assessment – enabled through acquisition of a much greater number of genome sequences from the sub-Antarctic and Antarctica – indicates a more complex dissemination scenario. The FIT phylogeographic analysis (Supplementary Figure S10; also Supplementary Figures S11-S14 for comparisons between the FIT and BIT phylogeographic modeling results) as performed in the previous publications suggested an initial spread from South Georgia to Crozet, and then from Crozet to Gough, Prince Edward and Kerguelen. This result was puzzling, as a virus transmission event from Crozet to Gough is unexpected given that at that time the mortality level at Crozet was very low. Moreover, most seabird species that are circumpolar migrants travel eastward (Figure 4). The issue of sampling bias here is relevant as Crozet is the only location where samples were collected in the winter and following a systematic carcass sampling protocol, regardless of any HPAIV suspicion. The BIT phylogeographic model explicitly accounts for the effects of viral migration events on the shape of the phylogeny, and is deemed more robust to the presence of such sampling bias than the FIT model ^53,54^. Our BIT phylogeographic analysis suggests initial and independent spread of Clade I from South Georgia to Gough, Crozet and Kerguelen, and from Crozet to Prince Edward. One single winter sample from Crozet is estimated to stem from a transmission event from Gough (Figure 3, Supplementary Figure S8), and hence constitutes a second introduction to Crozet, which did not lead to further transmission. Our BIT model parameterisation does not rely on any *a priori* knowledge of the relative distances and positioning of the islands, yet still provides a dispersal scenario that aligns with expectations based on seabird movements (Figure 4). However, unlike all other locations, there were no large-scale outbreaks detected on Gough. Hence it is unexpected that Gough would be a transmission hub or a stepping-stone for further transmission to Crozet (but see the Discussion section for a paragraph concerning at-sea transmission).

**Figure 4.**
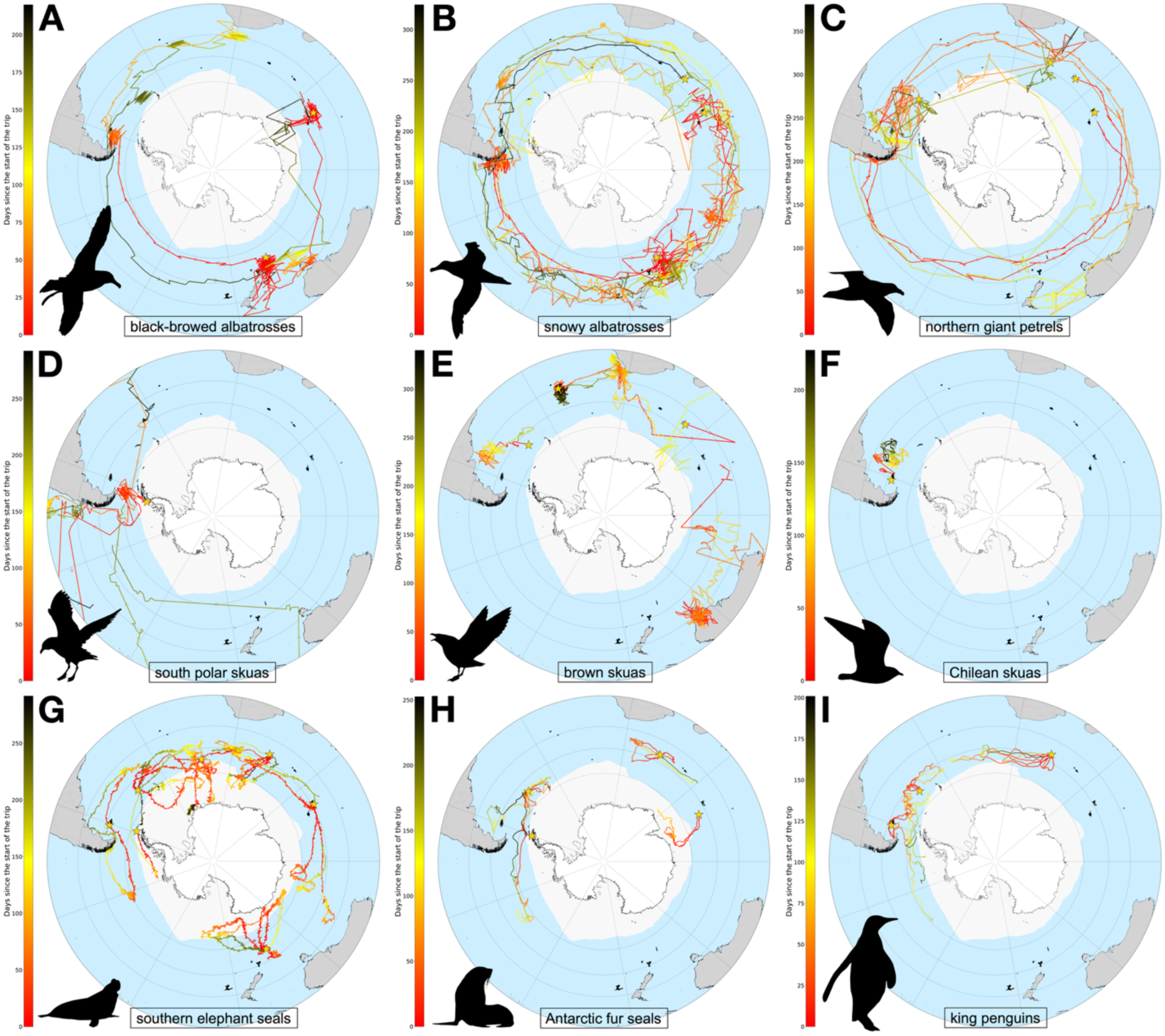
Selected tracking data for relevant species in the sub-Antarctic and Antarctica. The selection is made to depict representative long-distance movements. A) black-browed albatrosses ^70^; B) snowy albatrosses ^61^; C) northern giant petrels ^69^; D) south polar skuas; E) brown skuas (during the non-breeding period ^64,65^); F) Chilean skuas ^65^; G) southern elephant seals ^62^; H) Antarctic fur seals ^62^; and I) king penguins ^62^. Yellow stars represent tagging locations. Red-to-black color gradients indicate days since the start of the trips shown. The flight directions are indicated with arrowheads. A location-annotated map can be found in Supplementary Figure S15. Animal silhouettes courtesy of PhyloPic ^71^.

### Local reemergence of both Clade I and Clade II during the 2024/25 austral summer

At the end of the 2024 austral winter, we detected new HPAIV H5N1-related mortality as soon as the annual breeding season started in September 2024 on the Falklands and South Georgia. The first reports from the Antarctic Peninsula came in December 2024 ^18^. Our phylogenetic analyses revealed that Clade II viruses reemerged in the Falklands (affecting almost exclusively gentoo penguins), in South Georgia (affecting almost exclusively Antarctic fur seals and southern elephant seals), and in the Antarctic Peninsula (affecting skuas and crabeater seals). All these reemergences were from lineages that were already present locally at each of these three locations during the 2023/24 breeding season. Clade I also reemerged on the Antarctic Peninsula from a lineage that was already present locally at the end of the previous season. These reemergences from clades that were previously locally present suggest either (1) sustained transmission locally within the few animals that remain at the site throughout the winter, (2) persistence of infectious material in the environment, or (3) sustained transmission between animals that would have kept interacting within the same social groups while being away during the winter. On the Falklands, the gentoo penguins remain locally, continue coming ashore and are social in winter ^40^. Given that they were the single species that was affected both before and after the winter, it is possible that gentoo penguins could have sustained the viral transmission throughout the winter. Conversely, on the Antarctic Peninsula, all affected species migrate, hence one of the latter two scenarios seems more plausible. On South Georgia, the most affected species before and after the 2024 winter were the southern elephant and Antarctic fur seals that migrate in winter, but various other species remain ashore during that time. As a result, none of the three scenarios clearly stands out from the others.

Following these local reemergences and as happened in the previous season, Clade II spread at least two more times from South Georgia to the Antarctic Peninsula, either at the end of the 2023/24 breeding season or the beginning of the 2024/25 breeding season. Clade II also spread at least twice from the Falklands to South Georgia and also at least twice from the Falklands to the Antarctic Peninsula. Clade II is hence characterised – both in 2023/24 and 2024/25 – by numerous between-island transmission events in the Scotia Sea (South West Atlantic). No transmission events from the Antarctic Peninsula to other locations were identified, and there have been no additional dispersal events detected from South America in neither Clade I nor Clade II, which is in line with the absence of HPAI reports in wildlife from coastal Argentina and Uruguay since February 2024.

### Long-distance tracking data of large Procellariiformes align with inferred dispersal of Clade I

The presence of long-distance transmission events in Clade I led us to summarise known long-distance movements of selected species that were exposed to HPAIV H5N1, i.e. the most affected species according to our dataset, together with northern giant petrels (*Macronectes halli*) as they were highly exposed despite facing subsequent low mortality in comparison. These movements illustrate the connectivity between breeding sites in the Southern Ocean, as well as clear differences between species in terms of long-distance movement direction and velocity (Figure 4). Regardless of the distance and direction of their movements, flying seabirds travel considerably faster than seals and penguins (e.g., southern elephant seals ^60^ and snowy albatrosses ^61^).

Some species are mid-distance foragers: king penguins and Chilean skuas (*Stercorarius chilensis*) remain within hundreds of kilometres around their breeding sites, without overlapping much with individuals from other ocean basins (Figure 4F, 4I). Other species such as the southern elephant seals, the Antarctic fur seals and the black-browed albatrosses, are long-distance longitudinal travellers, and they can travel several thousand kilometers away from their breeding sites, mostly staying within the sub-Antarctic and Antarctic waters (Figure 4A, 4G, 4H); wintering areas of individuals from different ocean basins overlap ^60,62^. However, black-browed albatrosses from the Falklands, the only place where they were affected by HPAIV H5N1, are local foragers and do not commonly display such long-distance movements ^63^. Brown (*Stercorarius antarcticus*) and south polar (*Stercorarius maccormicki*) skuas are latitudinal migrants; they migrate north in winter, and wintering areas of individuals from distant ocean basins overlap (Figure 4D, 4E) ^64–66^. South polar skuas travel far further north, up to the northern hemisphere ^67,68^. Northern and southern (*Macronectes giganteus*) giant petrels (southern giant petrel tracks are not displayed in Figure 4 but are highly similar ^69^) and snowy albatrosses tend to remain within the same ocean basin as adults, but some individual adults, and all juveniles make longer-distance movements, including circumpolar (Figure 4B, 4C) ^69^; they can circumnavigate Antarctica in periods as short as 100 days ^61^.

Our phylogeographic reconstruction shows that the main direction of Clade I dispersal is eastward, which aligns with the dominant direction of long-distance movements of non-breeding large albatrosses and petrels which often include circumpolar movements, following the main wind direction (Figure 4). However, winter ranges of brown skuas and elephant seals from distant archipelagos overlap, and stepwise virus spread through transmission between those shorter-distance migrants is not impossible. Additionally, long-distance dispersal of the virus could also involve relay transmission between different host species. For example, after having been exposed on land, an elephant seal can potentially die at sea hundreds of kilometers away from its original colony. It may then be scavenged at sea by a seabird, potentially coming from another island hundreds of kilometers away. This example is important to illustrate a simple yet sensible transmission scenario that involves multiple species, but cannot be identified with our analyses since we did not sample carcasses at sea. We hence do not aim to identify a single species responsible for long-distance virus transport, but rather to show that such long-distance transmission events are possible given the movement patterns and may involve multiple hosts.

### HPAIV adaptation dynamics through lineage replacement and fixation of mutations at South Georgia

Our phylogeographic analysis shows that three HPAIV lineages succeeded each other on South Georgia. Each stemmed from distinct introduction events. We designate them in the following paragraph lineages A, B and C, by chronological order of case detection. Lineage A branches in Clade I, and was the only detected lineage between September 2023 and December 2023. By January 2024, lineage A was already almost entirely replaced by lineage B, which branches in Clade II (Figure 5). In the following breeding season (September 2024 to March 2025), not only had lineage A entirely disappeared, but lineage B - that accounted for almost the entirety of January 2024 cases - had also disappeared. Ultimately, only lineage C, also branching in Clade II, remained despite having been detected in only a single carcass in January 2024. Throughout the course of this succession of lineages, affected species were only seabirds at the very beginning, but seal mass mortality picked up quickly after (in November and December 2023). During the austral summer 2024/25, almost exclusively seals were affected, with a few individual snowy albatrosses being the only exceptions.

**Figure 5.**
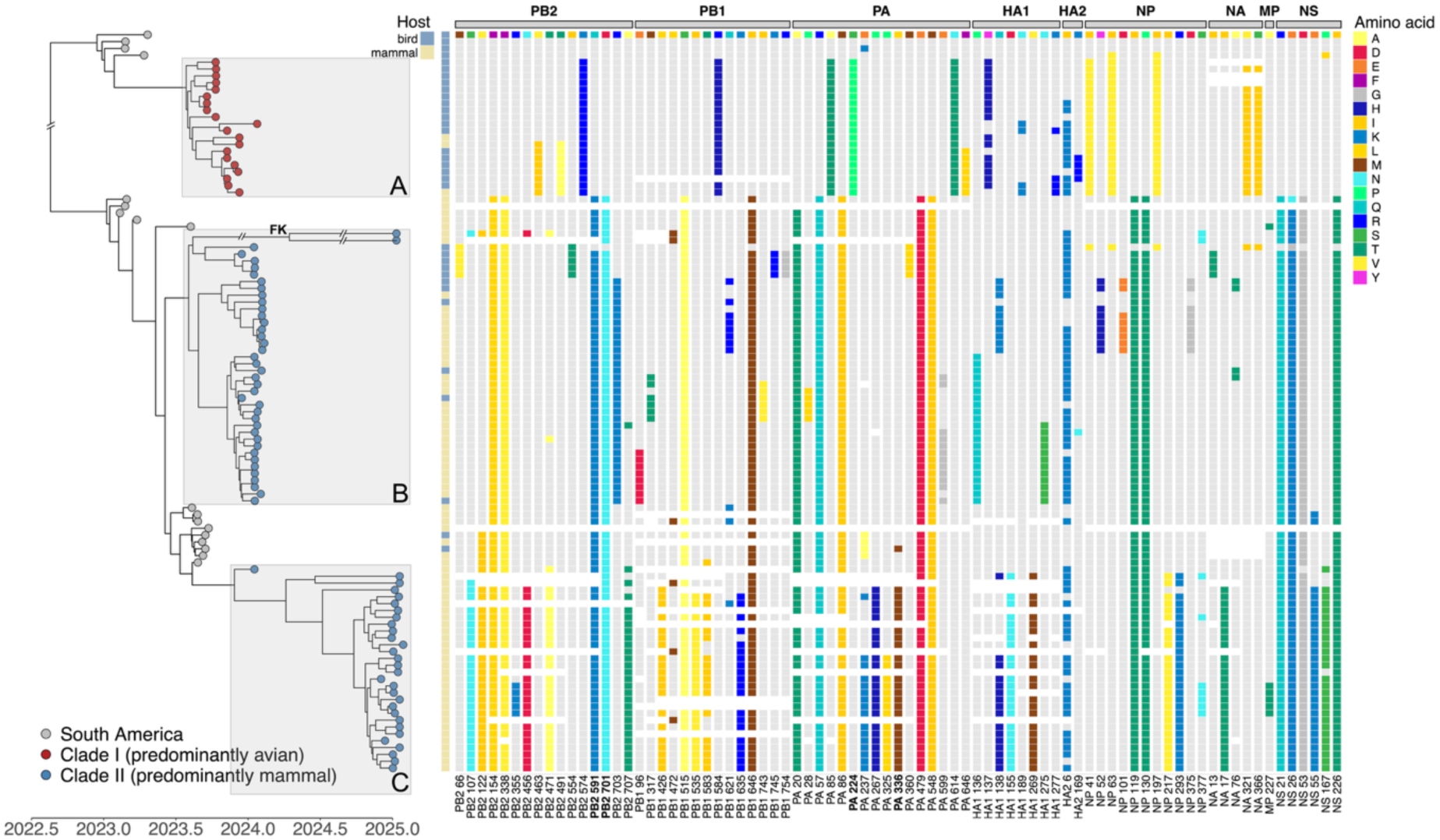
Overview of mutational differences between the predicted sequences of the proteins PB2, PB1, PA, HA1, HA2, NP, NA, M1 and NS1 coded by the HPAIV H5N1 2.3.4.4b genome obtained from South Georgia samples. Mutations with at least two occurrences in South Georgia sequences are depicted. The mutation indexing, at the bottom of the figure, is based on the mature HA. The sequences cluster into 3 different phylogenetic lineages, which we annotate according to chronological order lineages A, B and C. Each of these lineages descends from a separate introduction. The first one belongs to Clade I while the second and third belong to Clade II (Figure 2). With the exception of two sequences – each a separate introduction from the Falklands – all infections on South Georgia originated from South America (see Figure 3 and Supplementary Figure S9). Each of the three lineages fixed a set of known mammalian adaptations on South Georgia. Lineage A, that belongs to Clade I, carries the new mutation PA S224P, which can increase substantially the virulence for mammals when in combination with PA N383D ^74^ which is already present in the genetic background of our entire dataset. Lineage B carries the PB2 mutations D701N and Q591K. Lineage C, in addition to D701N and Q591K, carries PA L336M, and these three mutations are known to increase viral replication efficiency in mammalian cells ^75–77^.

These dynamics, with successive lineages replacing each other, suggest that the ongoing evolution of the viruses resulted in fitness differences not only between Clade I and Clade II, but also between different lineages within Clade II ^72,73^. Of note, later in the 2024/25 breeding season, two new introductions from the Falklands were detected (see Figure 5; annotated with ‘FK’) with the genetic composition of the second introduction event.

South Georgia lineages A, B and C are each defined by a set of nonsynonymous mutations that reached fixation – or near-fixation – at South Georgia, soon after the emergence of each of these lineages (Figure 5). We reviewed which of these mutations had previously been suggested to be related to host adaptation of influenza A viruses. Given that the vast majority of infected hosts at South Georgia were marine mammals, we expected to find mammalian adaptations. In lineage A, we identified the mutations PA S224P and NP I41V. PA S224P has been shown to substantially improve the replication efficacy in mammalian cells and the virulence for both mice and ducks when combined with the mutation PA N383D ^74,78^, which is present in all the sequences of our dataset – it was fixed long before the virus reached our regions of interest. The mutation NP I41V is known to enhance replication efficiency in mammalian cells at temperatures reproducing the conditions of the upper respiratory tract of mammals – lower than the temperature conditions in avian hosts, typically 41°C in their coelomic cavity ^79,80^. Lineage B is almost identical to the South American clade that was circulating among marine mammals ^11,58^. It has the mutations PB2 Q591K and PB2 D701N that were previously shown to have reached fixation in South America (Supplementary Figure S16). These two mutations are well-known to significantly increase the virus’ abilities to replicate in mammalian cells ^76,77^. It additionally has the mutations NS1 R21Q that also reached fixation in South America. NS1 residue 21 has been shown to be critical in impairing the antiviral response of the host cells via interaction with the retinoic acid-inducible gene I, which is host-specific and is therefore tightly linked to host adaptations – not specifically to mammalian adaptation ^81^. Lineage C shares all potential host adaptations that we identified in lineage B and that were already fixed in South America, but also newly acquired three additional ones – PA L336M, PA E237K and NS1 E55K. The mutation PA L336M is well documented and was present in the 2009-pandemic H1N1 virus – whose polymerase acidic protein was of avian origin – and has been shown to increase the polymerase efficiency in mammalian cells and the virulence of the virus for mice ^75^. PA residue 237 has also been shown to strongly influence the virulence of H5N1 viruses in ducks – 237K, the residue observed in the lineage C sequences, being associated with less virulence ^82^. NS1 K55E is not a well-established or high-impact mutation in H5N1, but has been associated with enhanced viral replication in human cells, mostly in synergy with NS1 K66E – already present in the genetic background of our entire dataset, and NS1 C133F, absent from our sequences^83^. Interestingly, the PB2 mutation E627K appeared once in lineage A and appeared several times at Crozet, and was detected in both avian and mammalian hosts (Supplementary Figure S16, Supplementary Figure S17). Since this mutation is well-known to be a critical mammalian adaptation ^46,84,85^, and since both on South Georgia and Crozet, mammalian mortality was intense, it is unexpected that this mutation did not reach a higher frequency.

On top of mutations related to host adaptation, we reviewed which mutations of the neuraminidase and of the hemagglutinin are located in known epitope sites and could be involved in antigenic drift. Remarkably, all mutations of the haemagglutinin that reached high frequency in any of the three lineages are located in regions of antigenic relevance. The mutations HA1 Y137H (lineage A), HA1 P136Q (lineage B), HA1 Q138K (lineage C) and HA1 S155N (lineage C) have not been specifically reported as sites under positive selection pressure, but are located within the receptor binding site in a region proposed to be of antigenic relevance ^86^. HA1 L269M is not located within the receptor binding site but has been found under positive selection pressure and suggested to be an antigenic site ^86,87^. None of the mutations of the neuraminidase that reached high frequency in any of the three lineages are located in known antigenic sites.

We also reported mutations that are known to affect the viral phenotype differences while not being directly linked to host adaptation and antigenic drifts. HA1 S155N is present in lineage C and is possibly not only involved in antigenic drift. It has been detected as an emerging mutation in US dairy cattle outbreaks, it is within the receptor binding site of the haemagglutinin and has been shown to increase binding to α2,3-linked sialic acid ^88^. NA V321I is fixed on lineage A and has been proposed to be a source of resistance to oseltamivir ^89^ antiviral treatment, however the efficiency of oseltamivir against lineage A has been specifically investigated and confirmed ^48^. The mutation HA2 I6K appeared recurrently on each of the three lineages, and approached fixation in each lineage. Given that this mutation is located inside the fusion peptide of the haemagglutinin, and that the substitution of an isoleucine – hydrophobic, by a lysine – polar and hydrophilic, is expected to drastically impair the fusion peptide function ^90^. The recurrence and frequency of this mutation in our sub-Antarctic and Antarctica sequences is unexpected and unexplained. The mutation HA0 A9T, fixed on lineage C, also is located within the signal peptide of the immature haemagglutinin and hence unexpected and we suspect it to be impactful ^91^. Besides these mutations, others have reached high frequency in at least one of the three lineages but we did not find in the literature anything about their phenotypic implications. In lineage A, this is the case of PB2 K574R, PB1 R584H, PA A85T, PA N614T, NP I63V, NP I197V, NA S366I. Only PB2 R703K is specific to lineage B, while lineage B and C share PB2 F154L, PB2 F338V, PB1 S515A, PB1 I646M, PA A20T, PA R57Q, PA M86I, PA E479D, PA M548I, NP I119T, NP P130T, NS1 E26K and NS1 D53G. Specific to lineage C, these mutations are: PB2 S107N, PB2 V122I, PB2 N456D, PB2 T471A, PB2 V707T, PB1 L426I, PB1 I535V, PB1 T583I, PB1 K635R, PA P267H, PA P325L, NP I217V, NP H293K, NA I17T and NS1 P167S.

## Discussion

Despite their remoteness, the ecosystems of Antarctica and the sub-Antarctic have been altered dramatically by human activities. Historically, extractive industries like whaling and sealing led to drastic depletions of the marine mammal biomass ^92^. The early human settlements associated with these industries additionally led to the accidental introduction of numerous non-native species, some of which turned invasive ^93^. This profoundly altered the ecosystems and a range of native seabird species have been extirpated from numerous locations due to predation by introduced species ^94,95^. More recently, bycatch and competition induced by the development of fisheries in Antarctica and the sub-Antarctic emerged as an additional threat ^96^. Climate change induces disproportionately higher temperature rises in polar regions than anywhere else on earth, placing increased pressure on the biodiversity and future viability of sub-Antarctic and Antarctic biodiversity ^95,96^. The recent arrival of HPAIV H5N1 2.3.4.4b in the region hence adds to the already long list of threats on these fragile biomes, and now constitutes a major species conservation concern in the region ^10^. Our results highlight the risk of persistence of the virus in the sub-Antarctic and in Antarctica. The implications could be disastrous if large-scale HPAIV H5N1 outbreaks become recurrent in these regions. The ongoing virus evolution and spread could lead to a larger number of species to be affected in the coming years. Mitigation measures are always difficult to implement – even more in these remote locations. However, recent king penguin vaccination trials on Crozet provide potential strategies to protect the most endangered species, provided the species and locations that are at risk can be determined in due time ^97^.

Building on earlier results based on smaller datasets ^14,17,19,20,48,49^, we demonstrate marked differences between the dispersal patterns of Clade I and Clade II within the sub-Antarctic and Antarctica, and suggest that the drivers of the transmission of each clade could differ significantly. On the one hand, Clade I spread tended to be longer-distance, eastward and circumpolar, to the South East Atlantic and South Indian Ocean. Remarkably, Clade I did not spread recurrently nor back and forth between locations, neither in the Atlantic nor in the Indian Ocean, and did not reach the Falklands. The long-distance spread of Clade I is consistent with the movements of several large Procellariiformes species, suggesting that these scavenging species could represent plausible vectors for the long distant spread of HPAIV ^98^. Given that these species circumnavigate Antarctica and breed throughout the region on sub-Antarctic islands and on the continent ^39^, they may act as long distance vectors of the virus. Consequently, the risk of Clade I spreading further east to other sub-Antarctic and Antarctica locations or to Oceania could be significant, as exemplified by the recent confirmation of HPAIV H5N1 on Heard Island ^99^, southeast of Kerguelen (Supplementary Figure S15).

On the other hand, Clade II remained within the Scotia Sea area where numerous and recurrent transmission events occurred between South Georgia and the Falklands and to the Antarctic Peninsula across both seasons. Almost all the species that were confirmed HPAIV H5N1-positive travel these distances within short time periods and could be plausible vectors for Clade II. Important exceptions are the crabeater seals, impacted on the Antarctic Peninsula and which are confined to the Antarctic region, and the gentoo penguins, impacted in the Falklands despite breeding at all sites – they remain close to their breeding site year round ^39^ and are hence unlikely involved in viral spread between archipelagos.

Furthermore, the three lineages introduced within the 2023/24 austral summer on South Georgia appeared engaged in local lineage replacement dynamics. The drastic, rapid and successive change of frequency of these lineages took place while the outbreak size remained high, hence suggesting adaptive dynamics rather than stochasticity alone ^72,73^. We found that each of the three lineages has a set of already known mammal-adaptive mutations, and hence that the succession of lineages at South Georgia provides an interesting opportunity to compare the relative fitness advantage of these different combinations of mutations. We also identified a set of undocumented mutations that reached fixation quickly - potentially simply due to selective sweep, but some might be worth further investigation.

The continued fixation of new mutations that are already known to increase the replication efficiency of influenza A viruses in human cells may suggest that the selection pressure on the polymerase complex of the virus is similar in seal and human hosts. This is especially concerning from a zoonotic perspective. Conversely, none of the observed mutations of the haemagglutinin are known to increase binding to α2,6-linked sialic acid, which is confirmed by a recent study investigating the sialic acid linkage of lineages A and B ^48^. Previous studies of influenza A virus binding in the respiratory tract of seals showed that human influenza viruses bind less efficiently than avian ones in the respiratory tract of seals ^100,101^ and α2,3-linked sialic acids have also been found predominant in the lungs of marine mammals ^102^. Our results support that transmission in seals likely does not benefit from increased binding to α2,6-linked sialic acid, hence the risk of emergence of such mutations – which would be worrying from a zoonotic perspective – may be limited. However, zoonotic viruses are often not pre-adapted ^103^, as supported by the numerous transmission events from birds to mammals and from mammals to birds that have been occurring since the first infections of marine mammals by HPAIV H5N1 in South America and now in the sub-Antarctic and Antarctica (Supplementary Figures S4 and S5)

At this stage it appears that neither of the two clades are mammalian or avian specific in the sub-Antarctic and Antarctica and both have been responsible for large numbers of deaths among birds and mammals. It comes as a surprise that on the Falklands, Clade II spreads almost exclusively among birds, and yet is retaining the mammalian adaptations (Supplementary Figure S17). This could suggest that those mammalian adaptations do not strongly reduce the ability of this clade to spread between birds. This is not dissimilar to the situation in South America, where Clade II initially disseminated primarily among pinnipeds but later also caused significant outbreaks in seabirds in Argentina ^58^. Of note, reverse mutation N701D was observed in a small cluster of terns there ^58^, while we have not observed it in albatross nor in penguin clusters in the Falklands. This could result from stochasticity only, but may suggest the D701N mutation is not equally deleterious in all avian hosts, such that the reverse mutation might not be strongly selected for in some avian hosts. A scenario in which highly mobile avian hosts disseminate viruses with mammalian-adapted mutations long-distance would be alarming, although to date Clade II has not spread nearly as far as Clade I.

Conversely, Clade I does not carry the PB2 mutations D701N nor Q591K, and despite its emergence, the mutation E627K has not reached a high prevalence. This is unexpected, given that Clade I has mostly been killing elephant seals at Crozet and Kerguelen, we would have expected these known mammalian adaptations to emerge and reach high frequency in HPAIV lineages spreading between mammalian hosts ^46,104^. This raises several non-exclusive hypotheses: the first one could be that other mutations, e.g. the PA mutations S224P and N383D ^74^, could compensate for the absence of PB2 mutations D701N and Q591K. A second hypothesis could be that spread between seals, unlike between other mammal species, might be efficient even in the absence of known mammalian adaptations, which would reduce the strength of the selective pressure in favour of these mutations. A third possibility could be that seals are only transmission dead-ends, but this would be unexpected given that elephant seals died even at locations where they are not mixed with birds. The specific case of the PB2 mutation E627K on Crozet leads us to another, fourth hypothesis. Its presence at intermediate frequencies might suggest that mammal-to-bird transmission events and vice versa are frequent, preventing the fixation of one phenotype or the other. As southern elephant seals and king penguins breed together in dense colonies on Crozet, this transmission scenario might be possible. These hypotheses trigger follow-up investigations of the relative sensitivity of seals compared to other mammals to influenza A strains that do not have mammalian adaptations (previous work already suggested they could be more sensitive ^105^), and serology surveys of the different species – including those that did not face high mortality – to unravel the finer details of the transmission dynamics.

### Limitations of the study

Our study does not provide estimates of the overall mortality across the sub-Antarctic and Antarctica. This is primarily due to the region’s remoteness and the complex logistics required to monitor isolated and widely distant sites. Exacerbating this, the tremendous number of carcasses make death counts of the most affected species and locations even harder. As an example, based on our field observations we think the Antarctic fur seals have, by far, suffered the highest number of deaths, but it has been impossible to enumerate this with any confidence. South Georgia hosts 98% of the world population of Antarctic fur seals, estimated to be over three million fur seals, with over one million pups born per year ^106^. The vast majority of sites visited on South Georgia had some level of HPAI-related mortality, with some sites already reaching over 50% of pup mortality weeks before the end of the 2024/25 outbreak (AC, JC, KO, KW, RN, JS unpublished data), but the mortality and carcass detection rates and carcass detection rates were substantially spatially heterogeneous. In such a context, estimating the total number of deaths is not possible with the currently available data. Another limitation of our analyses is that it relies almost exclusively on samples collected from carcasses. The host species represented in our dataset hence represent mostly the species that died from the disease – those that did not die from the viral infection are not represented. The phylogenetic analyses thus rely on the assumption that the host species representation in carcasses is representative of the infected host species.

Our sampling strategy relies on collecting samples from carcasses on land. While this is likely the only way to proceed, it cannot account for animals dying at sea. At-sea scavenging (Supplementary Figures S1-S2) may be an important transmission pathway, especially in winter when most species are at sea. We obtained sequences from snowy albatrosses, sooty albatrosses and Cape petrels, i.e., species that would not be exposed on land because they do not breed in dense colonies and do not scavenge on land. They could have been exposed to the virus by scavenging on floating carcasses at sea. Given the foraging range of these species, exposure could have happened hundreds to thousands of kilometres (Figure 4), away from the location where they were sampled, which our phylogeographic analyses does not account for.

Moreover, the impossibility to collect samples at sea implies that the winter transmissions are mostly unsampled. The sampling process is hence strongly biased. As a result, our phylogeographic reconstructions employed a BIT model ^53,54^ because it is less sensitive to sampling biases than the FIT model ^52^. However, we are ultimately unable to estimate the extent to which this is biasing our analyses and hence consider that the long branches of our consensus trees – which contain long unsampled ancestry – are to consider with caution.

## Methods

### Study areas

The study areas encompass all locations where HPAIV H5N1 infections were reported in the sub-Antarctic and Antarctica (Supplementary Figure S15). Because of their differences in size and accessibility, in some archipelagos not all islands were visited; on larger islands, only accessible sites were monitored. This led to important differences in monitoring between the different locations, in that on smaller islands HPAIV monitoring can be performed almost exhaustively, while on larger islands only the areas in proximity to landing sites and research stations were monitored. On multi-island archipelagos, HPAIV mortality has been assessed exclusively on the islands that were visited. In the Falklands (-51.698, -57,875) – the only inhabited sub-Antarctic islands reached by HPAIV H5N1 – most islands were visited and HPAI mortality was monitored all over the archipelago, based both on community reports and on scientific monitoring. On Crozet, HPAI mortality was monitored thoroughly on Possession Island (-46.433, 51.860; see Figure 2 in our previous study ^17^) but other islands were not visited. On Kerguelen (-49.352, 70.218), HPAIV H5N1 mortality was monitored in the areas accessible from the research station at Port-aux-Français, the eastern third of the island. Detected mortalities remained confined to a small portion of the southern coastline of the Courbet Peninsula (shown in more detail in Figure 2 of our previous study ^17^). Within Prince Edward, only Marion Island (-46.875, 37.859) was visited and thoroughly monitored for HPAI mortality. On Gough (-40.350139, -9.879), the monitoring was carried out at long-term monitoring sites across the island. On South Georgia (-54.283, -36.494), HPAIV H5N1-suspected cases are reported by cruise ships and by scientists based at the Bird Island and King Edward Point research stations. Hence, areas surrounding the research stations and cruise ship landing sites were regularly monitored. Samples were collected by scientists and were hence mostly collected in the monitored areas around the two research stations. However, this was complemented by scientific boat-based HPAIV H5N1 monitoring and sample collection in January 2024, November 2024 and January 2025, which provided opportunities to collect samples at sites where cruise ships reported mortality, and at additional sites that were not previously visited. On the Antarctic Peninsula (-63.462, -57.165) and neighboring Antarctic islands, HPAIV H5N1 mortality suspicions were reported by cruise ships, by scientists based on the various research stations, and by boat-based scientists. The monitoring and sample collection were discontinuous but rather thorough at a large number of sites that were visited.

### Sample collection

Brain, oropharyngeal and cloacal/rectal genetic samples were collected from fresh carcasses using sterile swabs and stored frozen, either without buffer, in lysis buffer (RNAlater (Qiagen), RNA/DNA Shield, eNAT System, InhibiSURE, etc.) or viral transport medium (VTM). In the Falklands, oropharyngeal swabs from live and healthy individuals were collected and stored in RNAlater.

### RNA extraction and sequencing

Samples collected on Prince Edward were extracted with an IndiMag Pathogen Kit (Indical Bioscience, Leipzig, Germany) with an IndiMag automated extraction instrument. Two amplification approaches were utilised, including a pan-genome RT-PCR assay ^107^ and leveraging the REPLI-g WGA & WTA Kit (Qiagen, Hilden Germany), using the whole transcription amplification protocol on half reaction volumes. The amplification products from both approaches were pooled and sequenced using an Ion Torrent at the Central Analytical Facility at Stellenbosch University, South Africa.

RNA extraction, whole genome amplification and sequencing of the samples collected at Crozet, Kerguelen and part of those of the Antarctic Peninsula and the Falklands was done using previously detailed procedures ^14,17,57^.

Another part of the Falklands and South Georgia sample set was processed at the Animal and Plant Health Agency, United Kingdom. RNA extraction was carried out using the MagMAX CORE Nucleic Acid Purification Kit (Thermo Fisher Scientific) as part of the robotic Mechanical Lysis Module (KingFisher Flex system; Life Technologies), according to the manufacturer’s instructions. Whole genome amplification and sequencing was then performed using previously detailed procedures ^14^.

For the last part of the samples from the Falklands and South Georgia, the RNA extraction was performed using the Zymo Quick-RNA™ Microprep Kit extraction kit and protocol, at the Department of Agriculture, Falkland Islands. The subsequent sequencing protocol consisted of a whole-genome amplification based on a previously published protocol ^107^. Amplified whole-genomes were then sequenced using MinION Nanopore sequencing, using the rapid barcoding kit and corresponding manufacturer instructions for amplicon sequencing.

### Phylogenetic analysis

Our starting data set for this study consisted of the 1221-taxa H5N1 clade 2.3.4.4b data set we constructed in previous work to study the circumpolar spread of H5N1 using a first batch of genomic sequences from the Crozet and Kerguelen Islands ^17^. To complement this dataset with recently uploaded sequences, on July the 12th, 2025, we downloaded all newly available H5N1 clade 2.3.4.4b sequence data from GenBank ^108^ and GISAID ^109,110^, focusing on Antarctica and South America sequences, i.e., where this lineage was spreading. We made sure none of the sequences from southern Africa were related to our lineage of interest. We complemented these data with those detailed in the sample collection section. We first aligned these sequences, and then added the resulting alignment to our original 1221-taxa alignment using MAFFT v7.490 ^111^, followed by manual / visual inspection, confirming that the collected data belong to the large HPAIV H5N1 2.3.4.4b clade that we previously analysed ^17^.

This yielded a data set containing 1383 concatenated viral genome sequences, with newly included sequences from the Antarctic Peninsula, Gough, Prince Edward, South Georgia, the Falklands, Crozet and Kerguelen. We first performed maximum-likelihood (ML) phylogenetic inference using IQ-TREE v3.0 ^112^ with automated model selection using ModelFinder ^113^, which selected the GTR+F+I+R6 model according to the Bayesian Information Criterion. We used this ML phylogeny as input for temporal signal assessment and outlier detection using TempEst ^114^. We did not detect any outliers in TempEst and hence our final data set consists of 1383 genomic sequences which exhibit a clear temporal signal.

We continued to perform a Bayesian time-calibrated phylogenetic analysis on this data set using Markov chain Monte Carlo as implemented in BEAST X (v1.10.5) ^115^, employing the BEAGLE high-performance computational library for computational efficiency to run on a powerful graphics processing unit ^116^. We employed a non-parametric coalescent model ^117,118^ as the tree prior, a general time-reversible substitution model ^119^ with a discretized gamma distribution to accommodate among-site rate heterogeneity ^120^, and an uncorrelated relaxed clock with an underlying lognormal distribution ^121^. We estimate the coalescent and clock model parameters using Hamiltonian Monte Carlo sampling ^122,123^. We used the default priors in BEAST X ^115^, including a conditional reference prior on the mean evolutionary rate ^124^. We ran this analysis until all relevant effective sample sizes reached at least 200, as assessed in Tracer v1.7.3 ^125^.

We checked whether independent replicate analyses in BEAST X converged to the same posterior distribution, both for the model parameters and for the phylogenetic trees ^126–128^. For the latter, we sampled 250 equally-spaced trees from each replicate’s posterior tree distribution and computed the pairwise Robinson-Foulds ^129^ (RF) and approximate subtree-prune-and-regraft (aSPR) ^130^ distances between all trees using R v.4.3.0 with the package *phangorn* ^131^. We plotted the coordinates for each combination of two of the first six dimensions^128^ from a classical multidimensional scaling analysis along with their topographical contour lines (not shown). Based on the RF and aSPR distances between phylogenies, we conclude that tree space for our phylogenetic analysis appears largely homogeneous and that the independent replicate analyses converge to the same posterior phylogenetic tree distribution^127^.

### Phylogeographic analysis

Based on the combined sampled posterior phylogenetic trees from this analysis, we performed a Bayesian discrete phylogeographic analysis using an asymmetric forward-in-time (FIT) continuous-time Markov chain model ^52^. The following locations were present in our 1383-taxa data set and used as discrete locations / states: Antarctica (incl. the South Shetland Islands), Argentina, Bolivia, Brazil, Canada, Chile, Colombia, Costa Rica, Crozet, Ecuador, Falklands, Gough, Guatemala, Honduras, Kerguelen, Prince Edward, Panama, Peru, South Georgia, United States, and Uruguay. We used the majority-rule highest independent posterior subtree reconstruction (HIPSTR) approach in TreeAnnotator X ^132^ to construct a time-calibrated consensus phylogeny, after removing an appropriate part of the chain as burn-in.

Given the presence of important temporal and geographic sampling bias in our comprehensive data set, we also performed two Bayesian discrete phylogeographic analyses – one on each identified clade of interest in the 1383-taxa phylogeny (Figure 2) due to computational restrictions – using a backward-in-time (BIT) structured coalescent approximation model (BASTA) ^53^, employing a recent implementation of the latter that markedly improves computational performance ^54^. Our first BIT phylogeographic analysis hence focused on the long-distance dispersal between South Georgia, Antarctica, Crozet, Kerguelen, Prince Edward and Gough. Our second BIT phylogeographic analysis focused on the (relatively) shorter-distance dispersal between South America, the Falklands, South Georgia, and the Antarctic Peninsula. We used the default priors in BEAST X ^115^, including a conditional reference prior on the geographic dispersal rate ^124^. To improve statistical mixing, we assume a shared population size between demes. We ran this analysis until all relevant effective sample sizes reached at least 200, as assessed in Tracer v1.7.3 ^125^. We again performed the aforementioned convergence and mixing assessments, and used the majority-rule HIPSTR approach in TreeAnnotator X ^132^ to construct a time-calibrated consensus phylogeny, after removing an appropriate part of the chain as burn-in.

### Non-synonymous mutation and recombination analysis

To identify all non-synonymous mutations, we aligned, phased and translated the putative proteins encoded by the genomic sequences of our dataset using VIGOR4 ^133^. We subsequently aligned and indexed the amino acid sequences. For the haemagglutinin, we used the H5 mature index, and we performed the variant calling on the amino acid alignment using a custom script with Python 3.

Additionally, we performed ancestral mutation reconstruction in BEAST X ^115^ on the PB2 Q591K, E627K and D701N amino acids using the complete genomes of the 628 sequences that make up the entire subtree containing Clade I and Clade II (Figure 2). We employed linked substitution, clock and tree prior models between a partition containing the nucleotide sequences of these three amino acids and a partition containing all other nucleotide positions. We performed a complete history reconstruction only on the amino acids of interest. We ran this analysis until all relevant effective sample sizes reached at least 200, as assessed in Tracer v1.7.3 ^125^.

We also performed single-gene phylogenetic analyses in BEAST X ^115^ to study signs of recombination, as shown in Figure 5. We focused on the PB2, NP and NA genes and employed the same approach as for the overall phylogenetic reconstruction on the full data set, but again restricted ourselves to the 628 sequences that make up the entire subtree containing Clade I and Clade II (Figure 2). We also ran this analysis until all relevant effective sample sizes reached at least 200, as assessed in Tracer v1.7.3 ^125^.

All associated figures were made using custom scripts with ggtree ^134^ in CRAN R ^135^.

### Movement tracking data of relevant species

To illustrate long-distance movement patterns of seabirds and seals across the Southern Ocean we present previously published and unpublished tracking data of the species that were the most affected by HPAIV H5N1, according to the available virus sequencing data, i.e., southern elephant seals, Antarctic fur seals, snowy albatrosses, king penguins, black-browed albatrosses, brown skuas, south polar skuas and Chilean skuas. Kelp gulls were not included because no tracking data from either the sub-Antarctic nor Antarctica are available. Gentoo penguins were not included because they are typically local foragers ^40^. Northern giant petrel tracking data were included because we observed them foraging on infected carcasses at numerous occasions (Supplementary Figures S1 and S2) and they might play an important role in the transmission of the virus, even though they did not face significant HPAIV-related mortality ^55^. Southern giant petrel tracks are not displayed because they are highly similar ^69^; both giant petrel species could play similar roles.

Tracking data for southern elephant seals, snowy albatrosses, king penguins, Antarctic fur seals and black-browed albatrosses were selected from the RAATD open access database ^62^ (accessed via GBIF.org on August 25th, 2025). The northern giant petrel data from Crozet are those previously published in Thiers et al. (2014) ^69^; those from South Georgia are unpublished data from 2017-2020 processed according to Merkel et al. (2016) ^136^, and those from Prince Edward are from the Seabird tracking database ^137^. The Chilean skua tracks from the Falklands and the brown skua tracks from South Georgia were published in Phillips et al. (2007) ^65^, those from Crozet in Delord et al. (2018) ^64^ and those from Gough in Steinfurth et al. ^59^. The south polar skua tracks are unpublished data from Rothera Point (Adelaide Island, Antarctic Peninsula) in 2017-2022, processed according to Merkel et al. (2016) ^136^. Individual tracks were manually selected to illustrate the type of long-distance movements for the different species. Hence, these movements do not aim to be representative of the average for their species, however they do not represent exceptional cases but relatively common movement types.

To build the background map, all coastlines except those of Antarctica and of the islands further south than 60°S were downloaded from the Global Self-consistent, Hierarchical, High-resolution Geography database ^138^ (accessed on July 31st, 2025). For visualisation purposes, the sizes of the smaller sub-Antarctic islands, i.e., Crozet, Amsterdam, Saint Paul, South Sandwich, Auckland, Campbell, Antipodes, Bounty, Snares, Bouvet, Gough, Tristan da Cunha, Chatham, Macquarie, Heard and the Prince Edward islands, are exaggerated by a six-fold factor, with the barycenter of each archipelago unchanged. The sizes of the bigger sub-Antarctic islands, i.e., the Falklands, South Georgia and Kerguelen are exaggerated by a 1.5-fold factor, also with the barycenter of each archipelago unchanged. The coastlines of Antarctica and of the islands south of 60°S were downloaded from the Scientific Committee on Antarctic Research geographic information system database ^139^. The country border lines are from the Natural Earth database (https://www.naturalearthdata.com/); we used the 110m resolution level. The sea ice extension line was downloaded from the National Oceanic and Atmospheric Administration database ^140^ (accessed on August 27th, 2025) and is the median extent of the sea ice front in September during the 1981-2010 period. The Antarctic Polar front, sub-Antarctic front and the subtropical front lines were drawn according to Orsi et al. (1995) ^141^ and downloaded from the Australian Antarctic division database ^142^. The projection is orthographic, centered on 10°E and truncated at 20°S.

The raw tracking data were processed through a custom cleaning pipeline that orders points in chronological order, and removes outliers, with outliers being defined as points that are too far out of the trajectories (distance superior to a species-specific threshold from the previous and next point). All maps were created using Python, using the *cartopy* ^143^, *shapely* ^144^, *geopandas* ^145^, *pandas* ^146^ and *matplotlib* ^147^ packages and their dependencies.

## Acknowledgements

The authors would like to thank the Department of Environment of the Falkland Islands Government ; the Department of Environment of the South Georgia and the South Sandwich Island Government, the Conservation Department of the Tristan da Cunha Government, and the French Southern Land authorities for granting permission to collect samples on the Falklands, South Georgia, Gough, Crozet and Kerguelen. The authors also want to thank all landowners in the Falklands that allowed us to collect samples on their lands.

The authors would like to thank the Department of Agriculture of the Falkland Islands government and especially Zoe Fowler and Josh Anderson-Wheatley for their support in the sample collection process and in the sample analyses and export, and for sharing labspace with us. We would also like to thank the Government of the South Georgia and South Sandwich Islands for their logistical support. We would like to thank the French Southern Land authorities for their technical support. We acknowledge the National Geographic and Rolex Perpetual Planet Ocean Expeditions and the Schmidt Ocean Institute aboard the R/V Falkor (Cruise ID FKt251214) for their technical support for sampling in the Weddell Sea. On Crozet and Kerguelen, we acknowledge critical technical support from the French Polar Institute (IPEV ECOPATH-1151, ORNITHO2E-109, ECOENERGY-119, ANTAVIA-137, ETHOTAAF-354, OIPLO-394, CYCLELEPH-1201, IA INTERPROJET-99). We thank Nicolas Keck, Gregory Jouvion, Rozenn Le Net, Karin Lemberger, Célia Lesage, Maxime Amy and SAGIR/Office Français de la Biodiversité (OFB) network for having contributed to organize the training of field personnel for wildlife disease surveillance which helped responding to HPAIV emergence on Crozet and Kerguelen Islands. We are grateful to Claudia Ulloa, Fabiola León, Carolina Márquez, Eduardo Pizarro, Lucas Krüger, Edna Correia, Rhiannon Gill, Vanessa Stephen, Peter Cunningham, Dylan Manyoka, Marcello Antonio, Megan Clarkson, Sibusisile Kheswa, Monique van Bers, Londani Rambua, Elmar van Rooyen, Karen Wolstenholme and Bernice Herwitt for their help with the sample collection. We would like to more specifically thank Jennifer Black for her important contribution to the sample collection in 2023-2024 at South Georgia.

Part of the sampling and sequencing on the Antarctic Peninsula was funded by ICM-ANID ICN2021_002 Millennium Institute BASE, ICN2021_044 Millennium Institute CGR, FONDECYT 1210568. On Kerguelen and Crozet, we also acknowledge specific support from Ceva Wildlife Research Fund since the beginning of the recent HPAI panzootic affecting wild species. The “NextSeq™ 2000” sequencing system and the “ThinkSystem SR650 V3 servers”, used to obtain and analyse the data at ANSES, were funded by a European Union grant and were used to sequence the Crozet, Kerguelen and part of the Falklands samples. For the Prince Edward islands, we are grateful to the ACAP Intersessional Group on High Pathogenicity Avian Influenza H5Nx for funding of sampling equipment and protective clothing, via ACAP’s Advisory Committee Small Grants Programme. The sequencing of Prince Edward samples was funded by the South African National Research Foundation under SARChI grant no. N00705. The sample collection at South Georgia was partly funded by the South Georgia and the South Sandwich Islands Government. Part of the testing and generation of the viral sequences from South Georgia, the Falklands and the Antarctic Peninsula was funded by the Department for Environment, Food and Rural Affairs (Defra, UK) and the Devolved Administrations of Scotland and Wales, through the following programmes: SV3400, SV3032, SV3006, SE2213 and SE2227. This work was also supported by the Biotechnology and Biological Sciences Research Council (BBSRC) and Department for Environment, Food and Rural Affairs (Defra, UK) research initiative ‘FluTrailMap’ [grant numbers BB/Y007271/1, BB/Y007298/1] and the Medical Research Council (MRC) and Defra research initiative ‘FluTrailMap-One Health’ [grant number MR/Y03368X/1]. Funded by the European Union under grant agreement (101084171) – (Kappa-Flu). Funding was also provided from Innovate UK (grant number 10085195). The rest of the samples from South Georgia and the Falklands was funded by Kenneth C. Griffin and Griffin Catalyst. The sample collection on Crozet and Kerguelen was funded by the French Polar Institute, by the ANR AAPG ECOPATHS project (ANR-21-CE35-0016), WILDFLU (ANR-25-CE35-0691)., REMOVE_DISEASE project (ANR-21-BIRE-0006; Biodiversa+ and Water JPI joint call for projects under the BiodivRestore ERA-NET Cofund GA N°101003777) and CNRS Ecologie & Environnement SEE-Life and HPAI initiative.

This article is a contribution to the Ecosystems component of the British Antarctic Survey Polar Science for a Sustainable Planet program, funded by the Natural Environment Research Council. PC acknowledge further support from FCT was received FCT through UID/04292 – Centro de Ciências do Mar e do Ambiente (MARE) and through the project LA/P/0069/2020 granted to the Associate Laboratory ARNET and from the Environmental Studies Budget of the Falkland Islands Government. For the bioinformatic analyses, M.B., S.L.H. and G.B. acknowledge support from the DURABLE EU4Health project 02/2023-01/2027 which is co-funded by the European Union (call EU4H-2021-PJ4) under Grant Agreement No. 101102733. Y.S. and M.A.S. acknowledge support from US National Institutes of Health grant R01AI153044. CPR acknowledges the support of the Laboratory of Molecular Virology UC Team and the grants INACH RT-30-22 and Centers of Excellence for Influenza Research and Response (CEIRR) Contract No. 75N93021C00017 Option 18A and 75N93021C00014. The WHO Collaborating Centre for Reference and Research is supported by the Australian Department of Health, Disability and Ageing. RET Vanstreels and MM Uhart were partially supported by the U.S. National Science Foundation Center for Pandemic Insights (NSF CPI), award number 2412522. This research was supported in part by the Intramural Research Program of the National Institutes of Health (NIH). The contributions of the NIH author(s) are considered Works of the United States Government. The findings and conclusions presented in this paper are those of the author(s) and do not necessarily reflect the views of the NIH or the U.S. Department of Health and Human Services, or the United States government. G.B. acknowledges support from the Research Foundation – Flanders (“Fonds voor Wetenschappelijk Onderzoek – Vlaanderen,” G098321N), and from the European Union Horizon 2023 RIA project LEAPS (grant agreement no. 101094685). Views and opinions expressed are however those of the author(s) only and do not necessarily reflect those of the European Union or REA. Neither the European Union nor the granting authority can be held responsible for them.

## Supplementary Materials

**Supplementary Figure S1.**
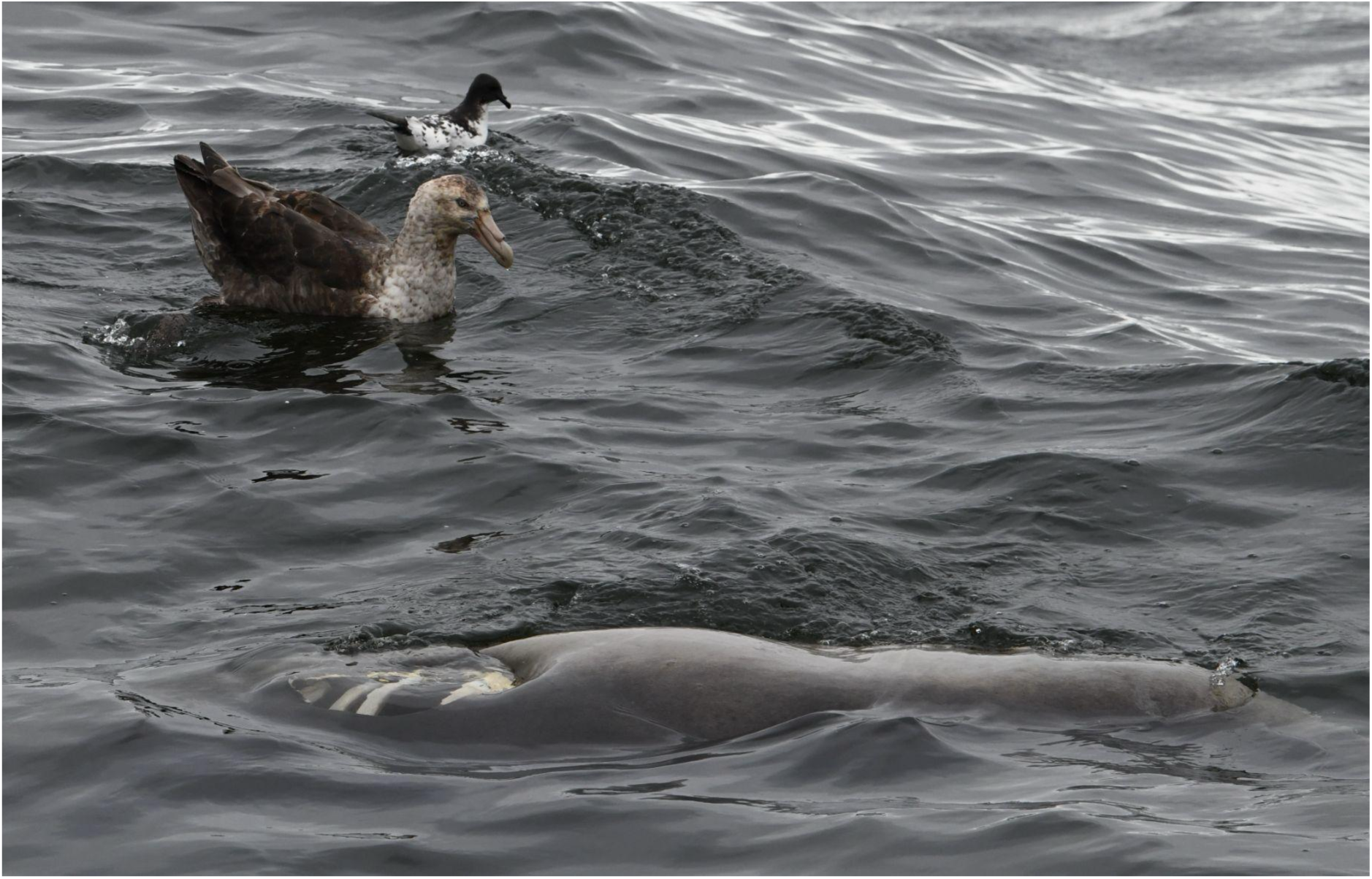
A southern giant petrel and a Cape petrel scavenging a dead southern elephant seal at sea, off the coast of South Georgia, on the 6th of December, 2025. Picture credit: Augustin Clessin.

**Supplementary Figure S2.**
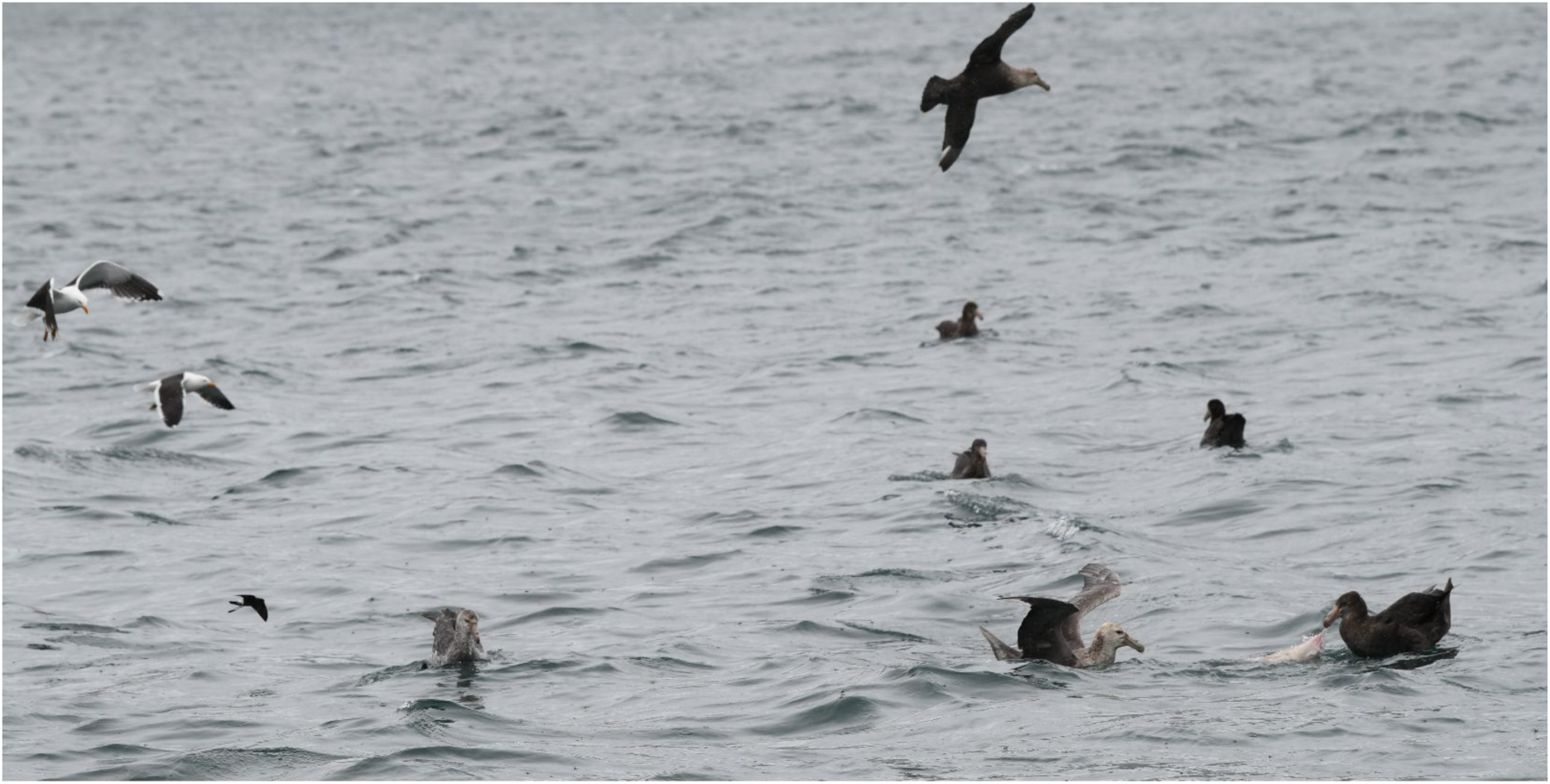
Southern and northern giant petrels, kelp gulls (*Larus dominicanus*) and a Wilson’s storm petrel (*Oceanites oceanicus*) scavenging an unidentified animal at sea, off the coast of South Georgia, on the 13th of December, 2025. Picture credit: Augustin Clessin.

**Supplementary Figure S3.**
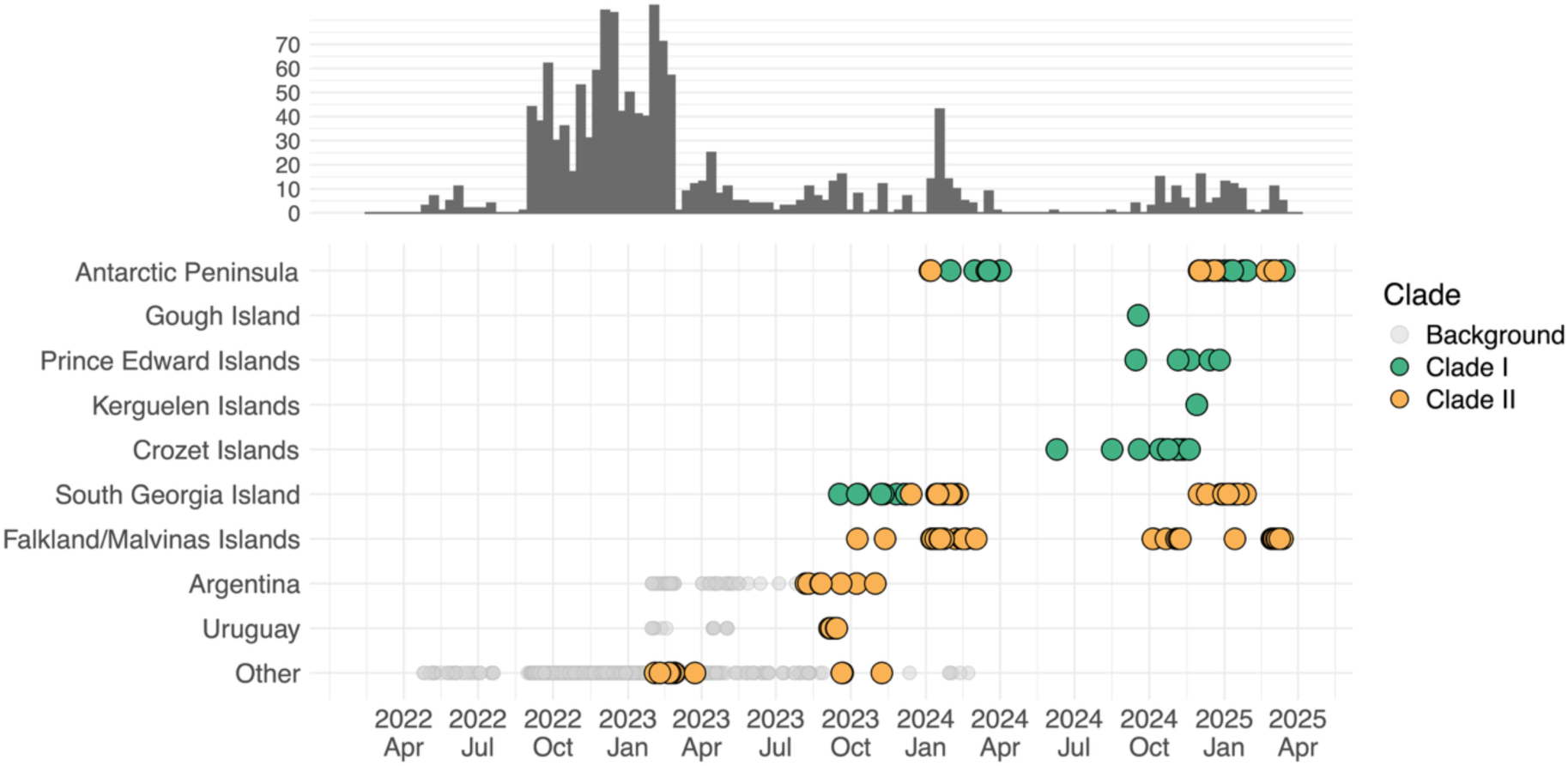
Known locations and dates in our well-supported phylogeny of genomes that contains the samples from the Antarctic Peninsula and the southern Indian Ocean. Color scheme focuses on genomes being part of Clade I, Clade II, or neither (i.e., background). Rows show the collection dates of genomes on the bottom, as well as the frequency of genomes as a bar plot at the top. Histogram bars use 2-week bins.

**Supplementary Figure S4.**
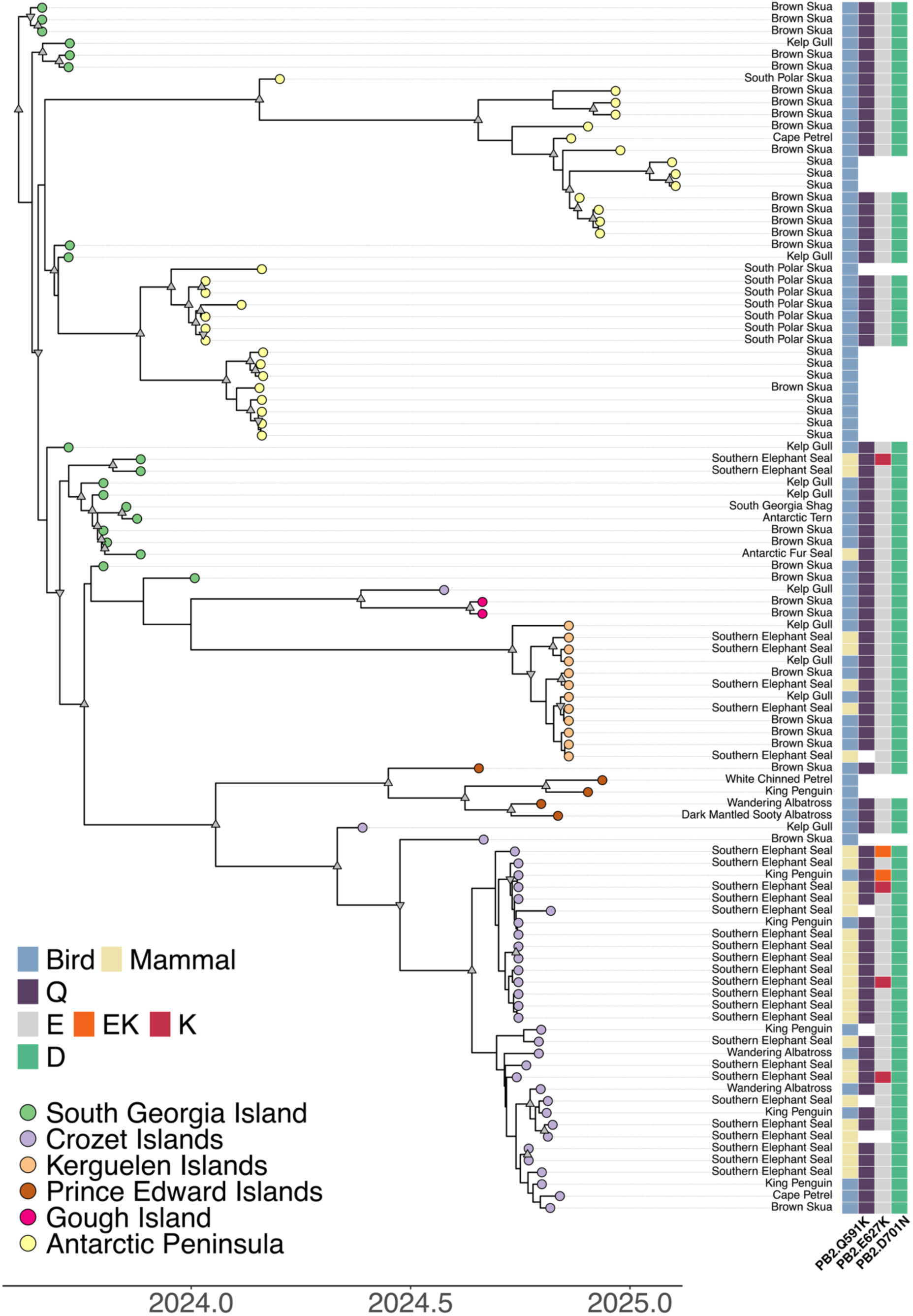
Time-calibrated majority-rule HIPSTR phylogeny of Clade I. Tip colors correspond to sampling locations, and internal nodes with posterior support above 0.7 and 0.9 are marked with down-facing and up-facing triangles respectively. Sampling host and the corresponding mammal or bird classification are shown for each taxon on the right.

**Supplementary Figure S5.**
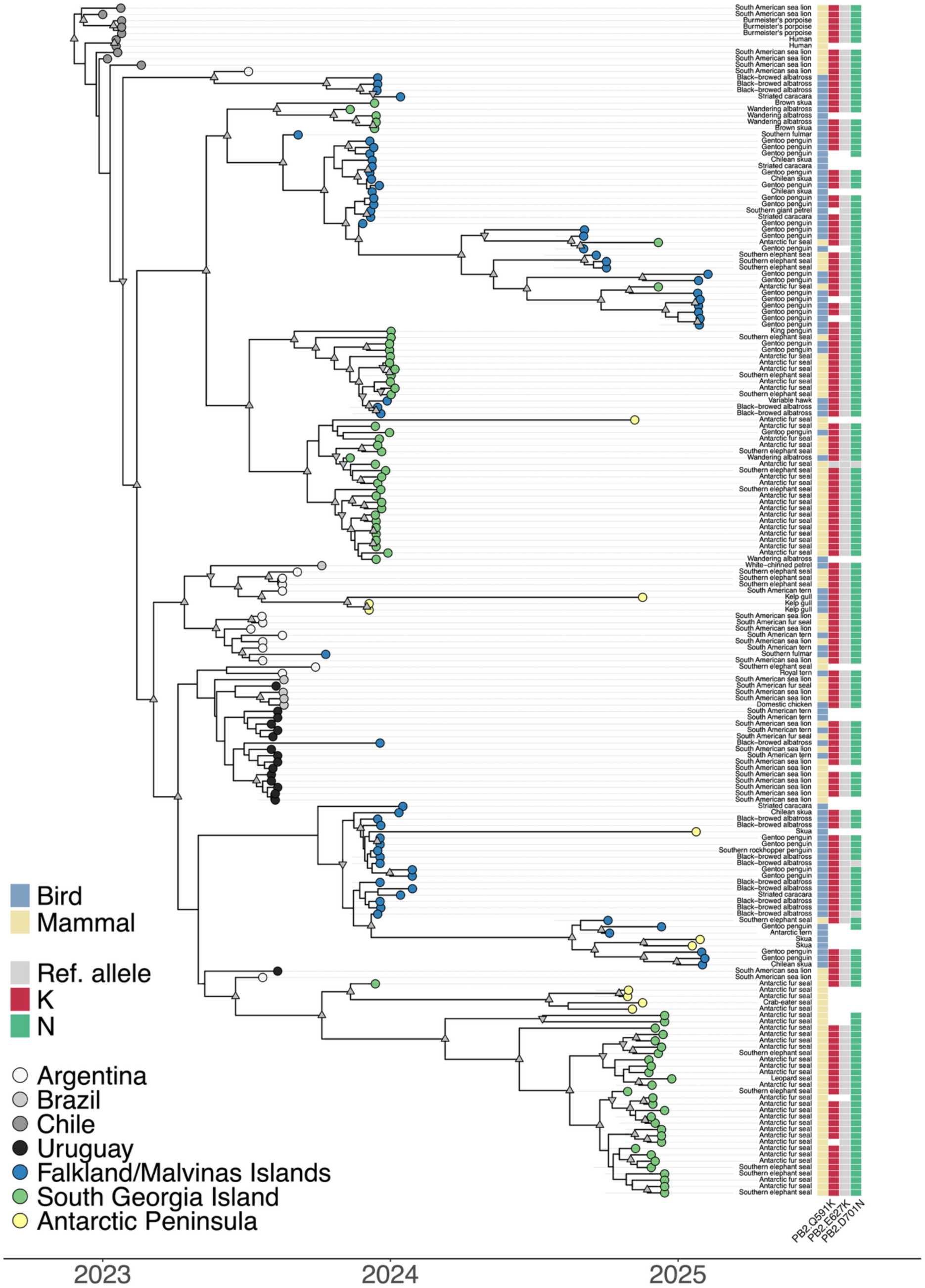
Time-calibrated majority-rule HIPSTR phylogeny of Clade II. Tip colors correspond to sampling locations, and internal nodes with posterior support above 0.7 and 0.9 are marked with down-facing and up-facing triangles respectively. Sampling host and the corresponding mammal or bird classification are shown for each taxon on the right.

**Supplementary Figure S6.**
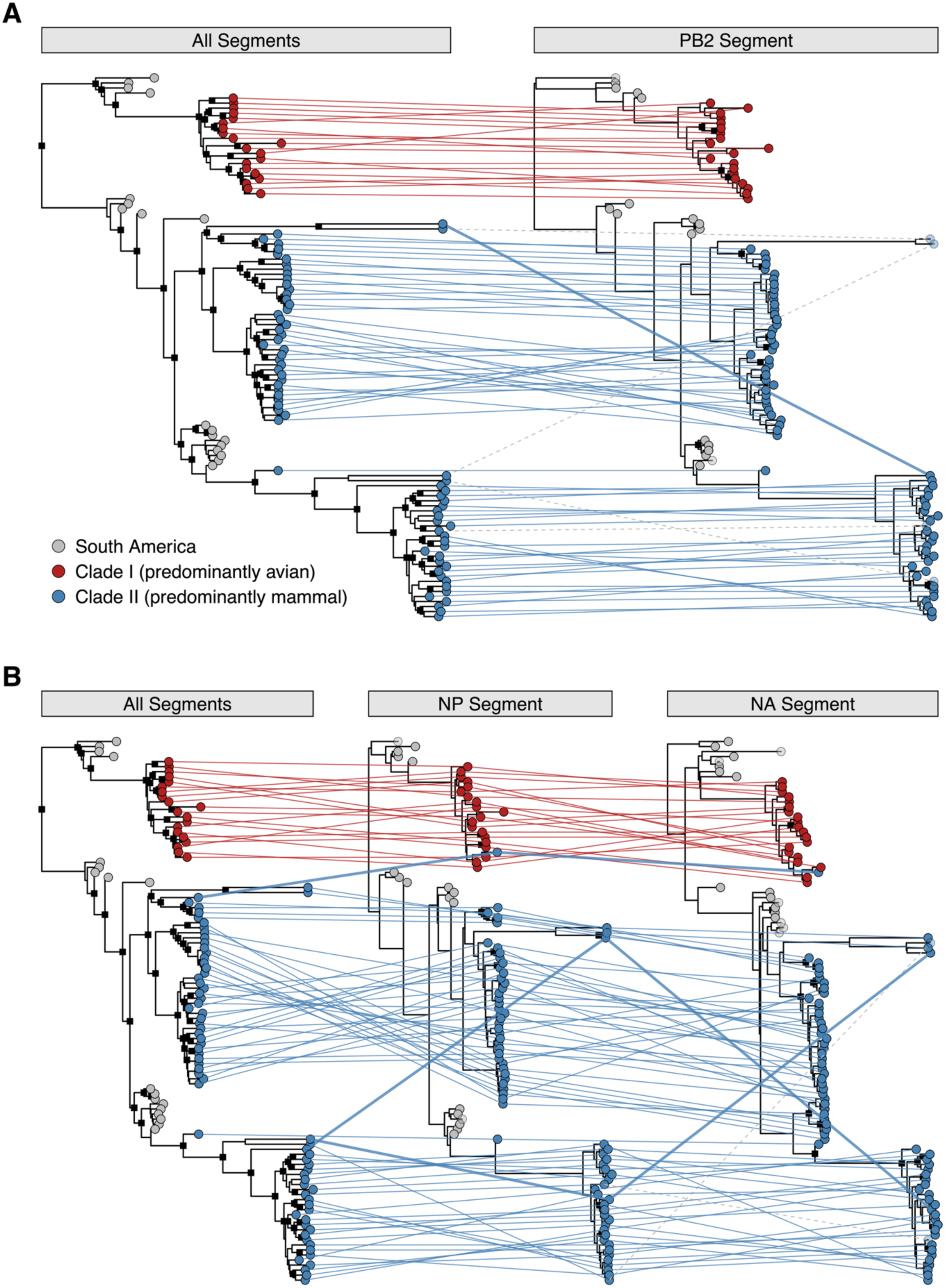
Tanglegrams identifying putative reassortment sequences by comparing the full genome phylogeny connecting the South Georgia lineages (see Figure 5) with those of the PB2, NP and NA genes. (A) One recombinant sequence with PB2 came from a sample taken on South Georgia on January 11th, 2025, from an Antarctic fur seal. (B) The top putative reassortment sequence came from a sample taken on Prior Island, South Georgia, on January 16th, 2024, from a brown skua; the middle putative reassortment sequence came from South Georgia on January 19th, 2025, from an Antarctic fur seal; the bottom putative reassortment sequence came from South Georgia on January 7th, 2025, from an Antarctic fur seal. Putative reassortments are connected with bold lines. Tips with less than 50% coverage are indicated with transparent circles, and dashed lines indicate the corresponding lines arriving in these low-coverage sequences (from which we hence draw no conclusions).

**Supplementary Figure S7.**
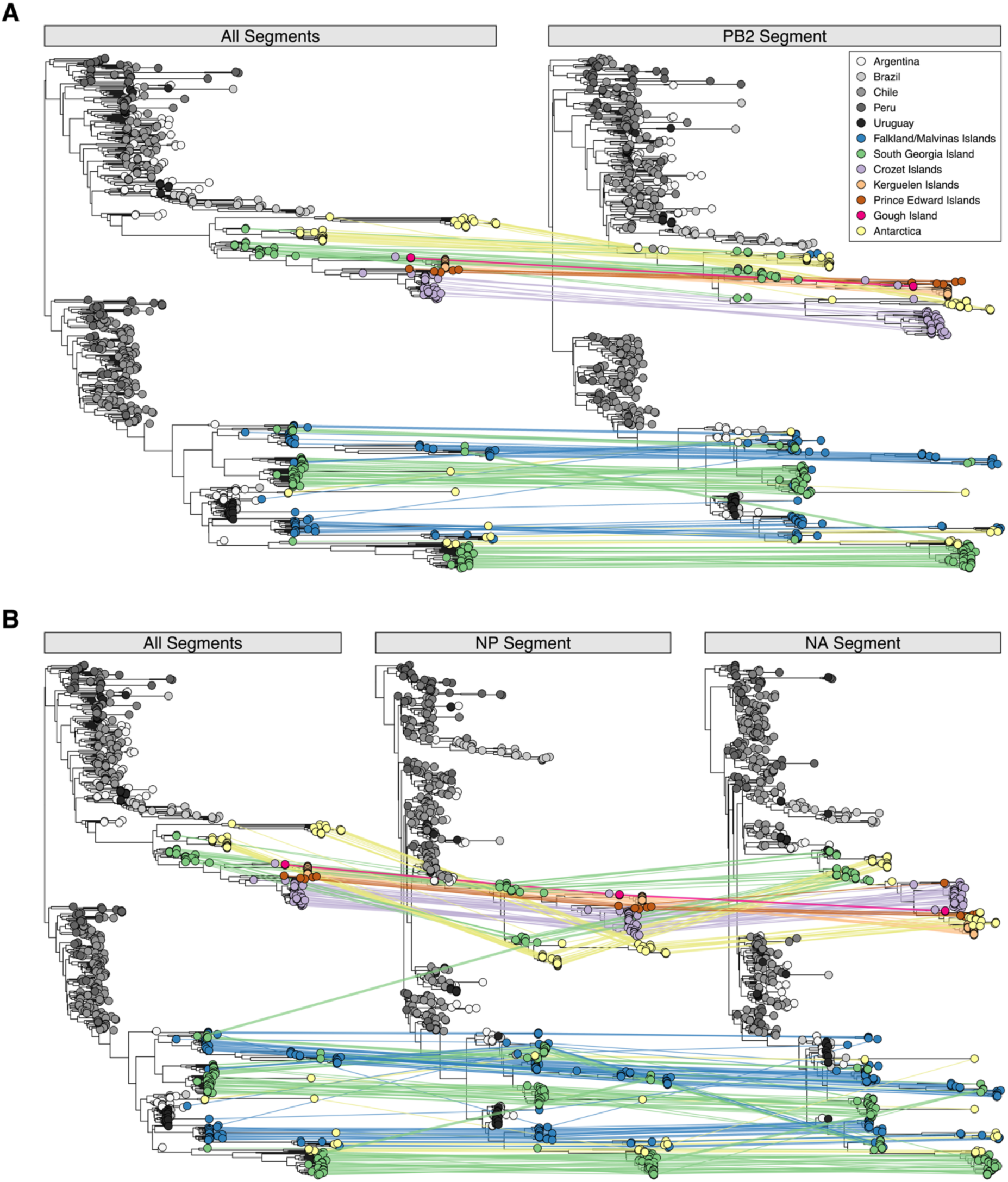
Tanglegrams identifying putative reassortment sequences by comparing the full genome phylogeny of the large subtree linking Clades I & II (see Figure 5) with that of the PB2, NP and NA genes. (A) The recombinant sequence came from a sample taken on South Georgia on January 11th, 2025, from an Antarctic fur seal. (B) The top putative reassortment sequence came from a sample taken on Prior Island, South Georgia, on January 16th, 2024, from a brown skua; the middle putative reassortment sequence came from South Georgia on January 19th, 2025, from an Antarctic fur seal; the bottom putative reassortment sequence came from South Georgia on January 7th, 2025, from an Antarctic fur seal. Putative reassortments are connected with bold lines. Tips with less than 50% coverage are indicated with transparent circles, and dashed lines indicate the corresponding lines arriving in these low-coverage sequences (from which we hence draw no conclusions).

**Supplementary Figure S8.**
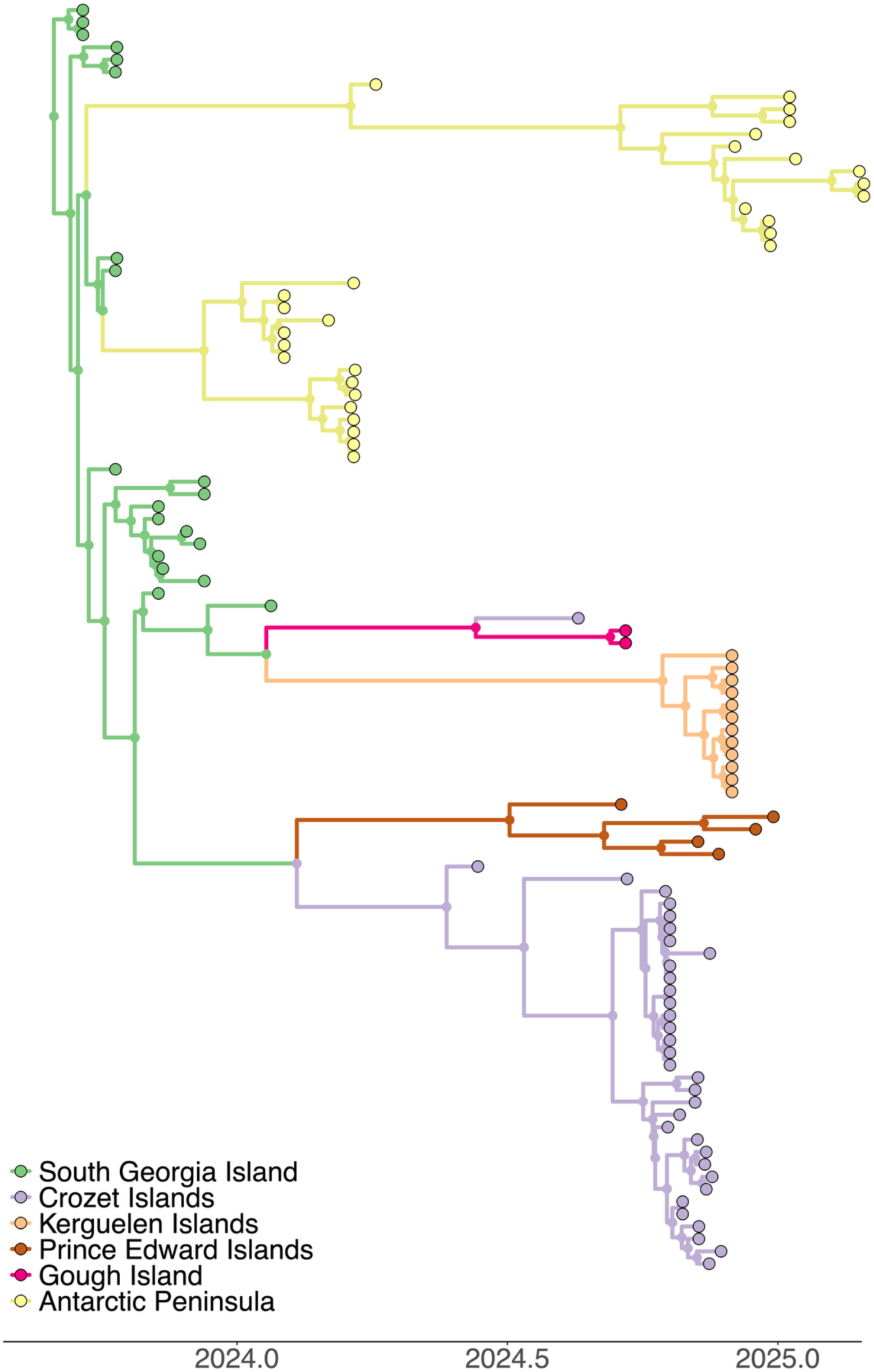
Clade I ancestral location inference using backward-in-time (BIT) phylogeographic inference. Two introductions into the Antarctic Peninsula are estimated to have occurred from South Georgia Island, which was also estimated to be the source of introductions into Crozet, Kerguelen, Prince Edward and Gough.

**Supplementary Figure S9.**
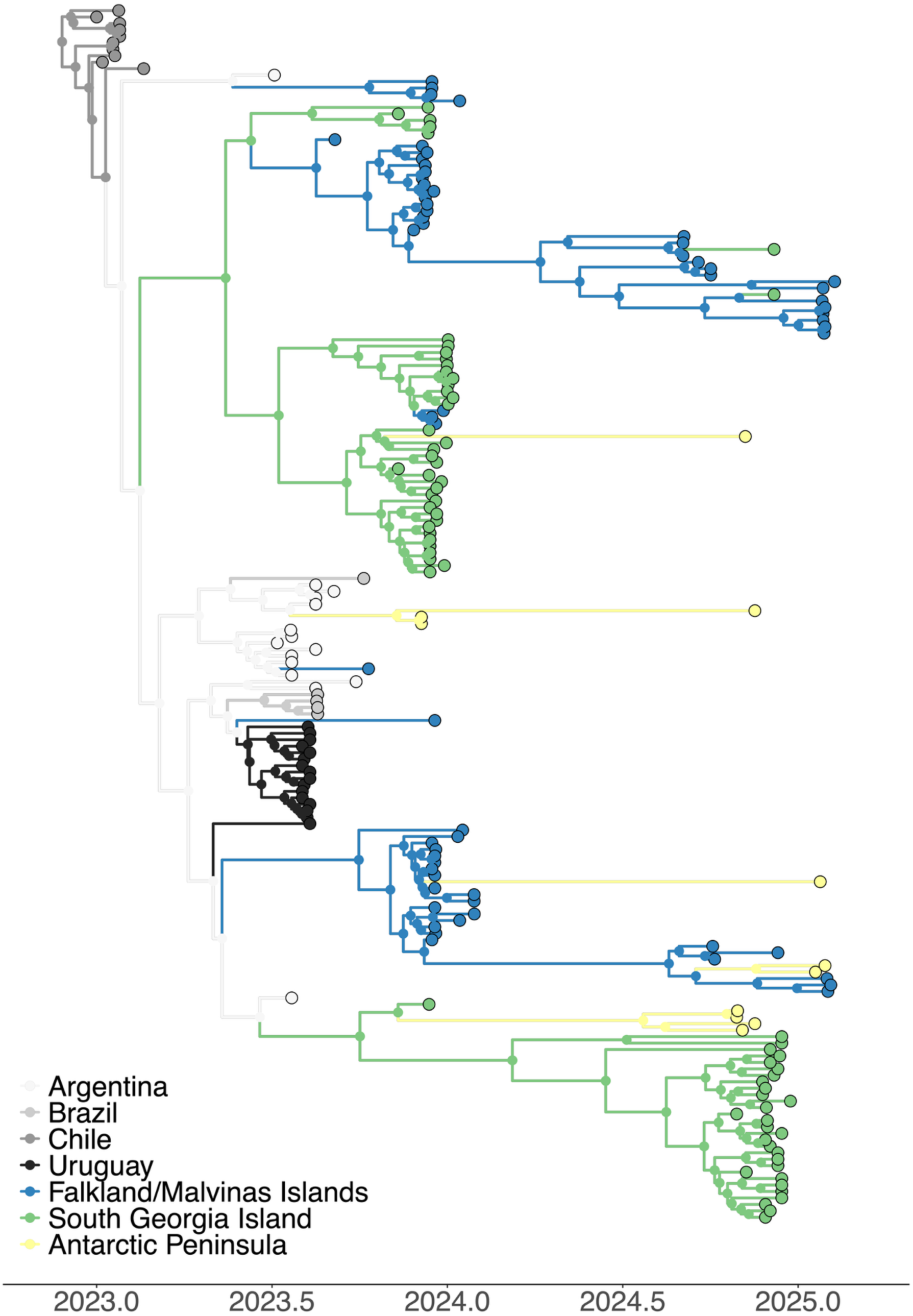
Clade II ancestral location inference using backward-in-time (BIT) phylogeographic inference. Introductions into the Falklands are estimated to have occurred from Argentina and South Georgia, which had its introductions originate in Argentina. The Antarctic Peninsula saw multiple introductions – leading to long branch lengths – from South Georgia, the Falklands and Argentina.

**Supplementary Figure S10.**
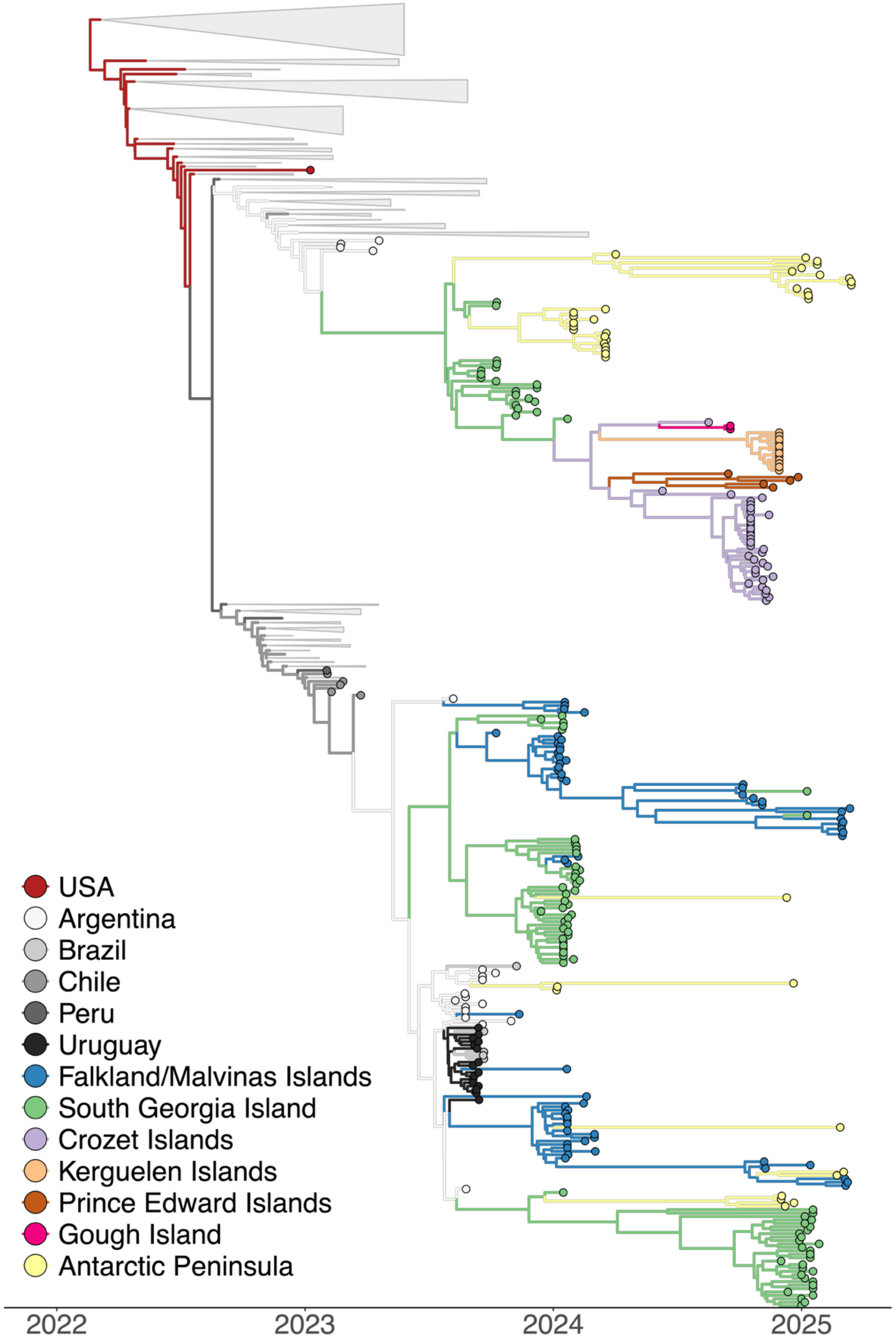
Origin and subsequent spread of HPAI H5N1 2.3.4.4b infections in the Southern Ocean according to a forward-in-time (FIT) discrete phylogeography model ^52^. Time-calibrated location-annotated phylogeny based on whole concatenated genomes, showing all inferred ancestral locations, shows a key difference with the BIT analysis (see Figure 3 and Supplementary Figure S6) in that the FIT model estimates a westward transmission event from Crozet to Gough, which is in contradiction with the absence of mass mortality at that time on Crozet.

**Supplementary Figure S11.**
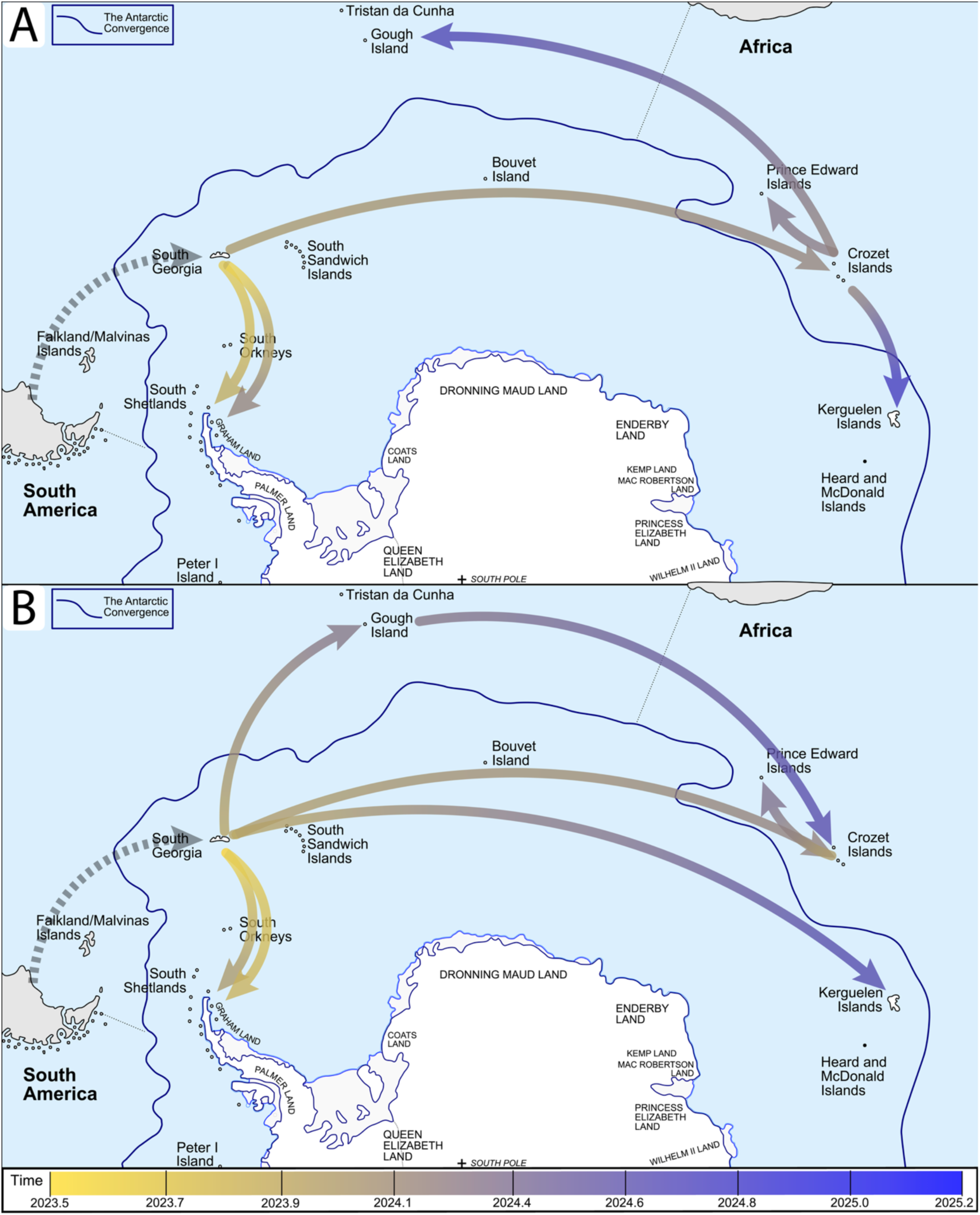
Improved phylogeographic modelling elucidates the long-distance circumpolar HPAI H5N1 2.3.4.4b dispersal towards Gough, Crozet, Kerguelen, and Prince Edward Islands (Clade I) after an initial introduction from South America into South Georgia (in grey; before the start of our time line). A) Forward-in-time (FIT) phylogeographic reconstruction ^52^ estimates an eastward dispersal from South Georgia to Crozet, and subsequently a westward dispersal from Crozet into Gough Island; B) Backward-in-time phylogeographic reconstruction ^53,54^ on the other hand estimates a consistent eastward movement. After an initial independent dispersal from South Georgia to Gough, Crozet and Kerguelen, the virus spread from Gough to Crozet and from Crozet to Prince Edward. The mass mortality of Crozet descends from the first introduction event, i.e., the one from South Georgia and not the one from Gough, that did not lead to further transmission. Arrows shown represent estimated single introductions and are coloured according to a time-dependent gradient. Background map adapted from ©Hogweard (talk · contribs), CC BY-SA 3.0 <https://creativecommons.org/licenses/by-sa/3.0>, via Wikimedia Commons.

**Supplementary Figure S12.**
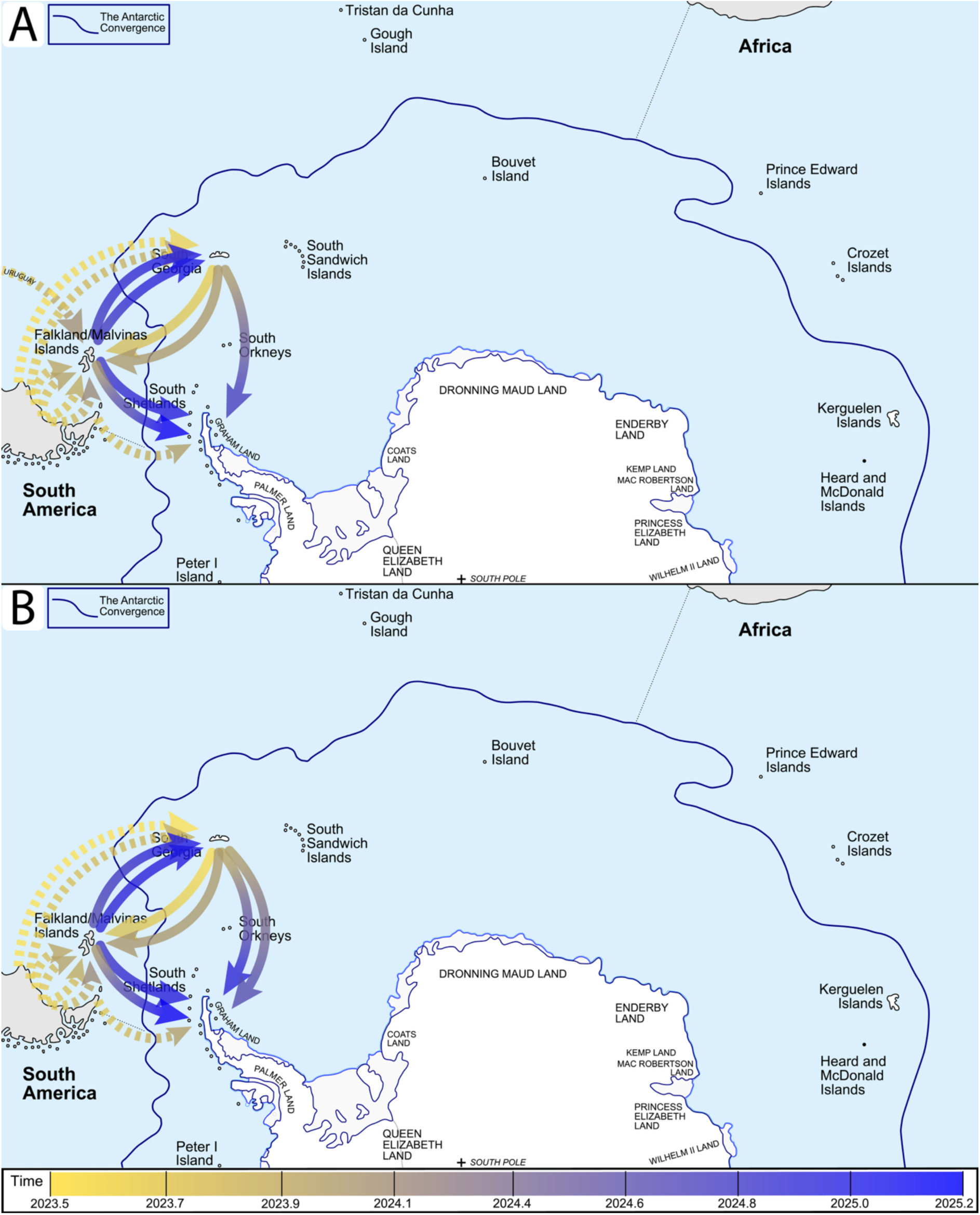
Phylogeographic modelling alternatives lead to similar estimates of the dispersal of HPAI H5N1 2.3.4.4b between the Falklands, South Georgia and Antarctica (Clade II). A) Forward-in-time phylogeographic reconstruction ^52^ estimates an additional introduction event from South America (Uruguay) into the Falklands and from South Georgia into the Antarctic Peninsula, compared to B) backward-in-time phylogeographic reconstruction ^53,54^, based on currently available data. Background map adapted from ©Hogweard (talk · contribs), CC BY-SA 3.0 <https://creativecommons.org/licenses/by-sa/3.0>, via Wikimedia Common

**Supplementary Figure S13.**
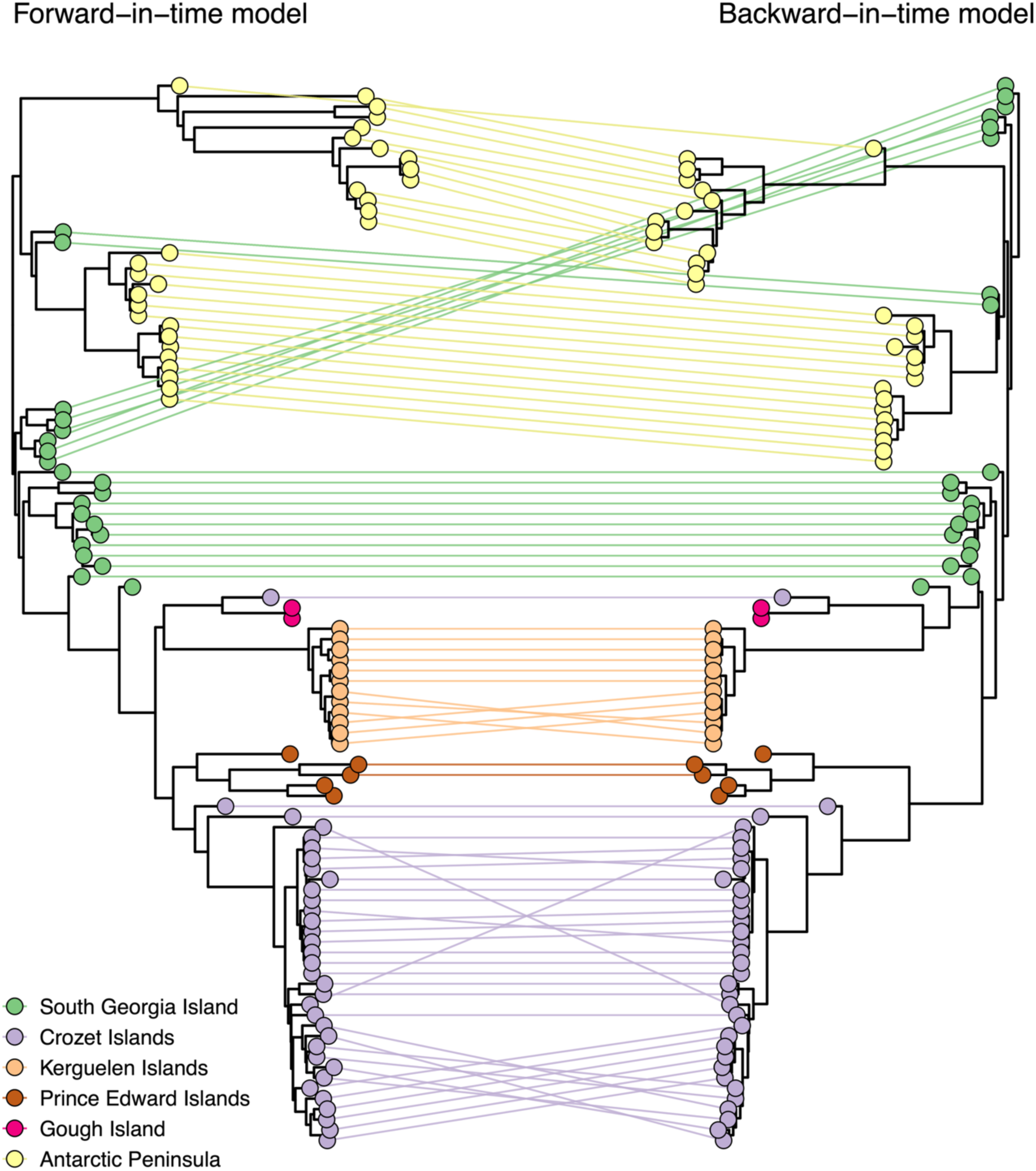
Tanglegram contrasting the majority-rule HIPSTR topologies of Clade I (see Figure 2) under the FIT (left) and BIT (right) phylogeographic models. Apart from minor within-location clustering differences, we observe a larger difference in the clustering of sequences from South Georgia which however does not have implications for estimated introductions into the Antarctic Peninsula.

**Supplementary Figure S14.**
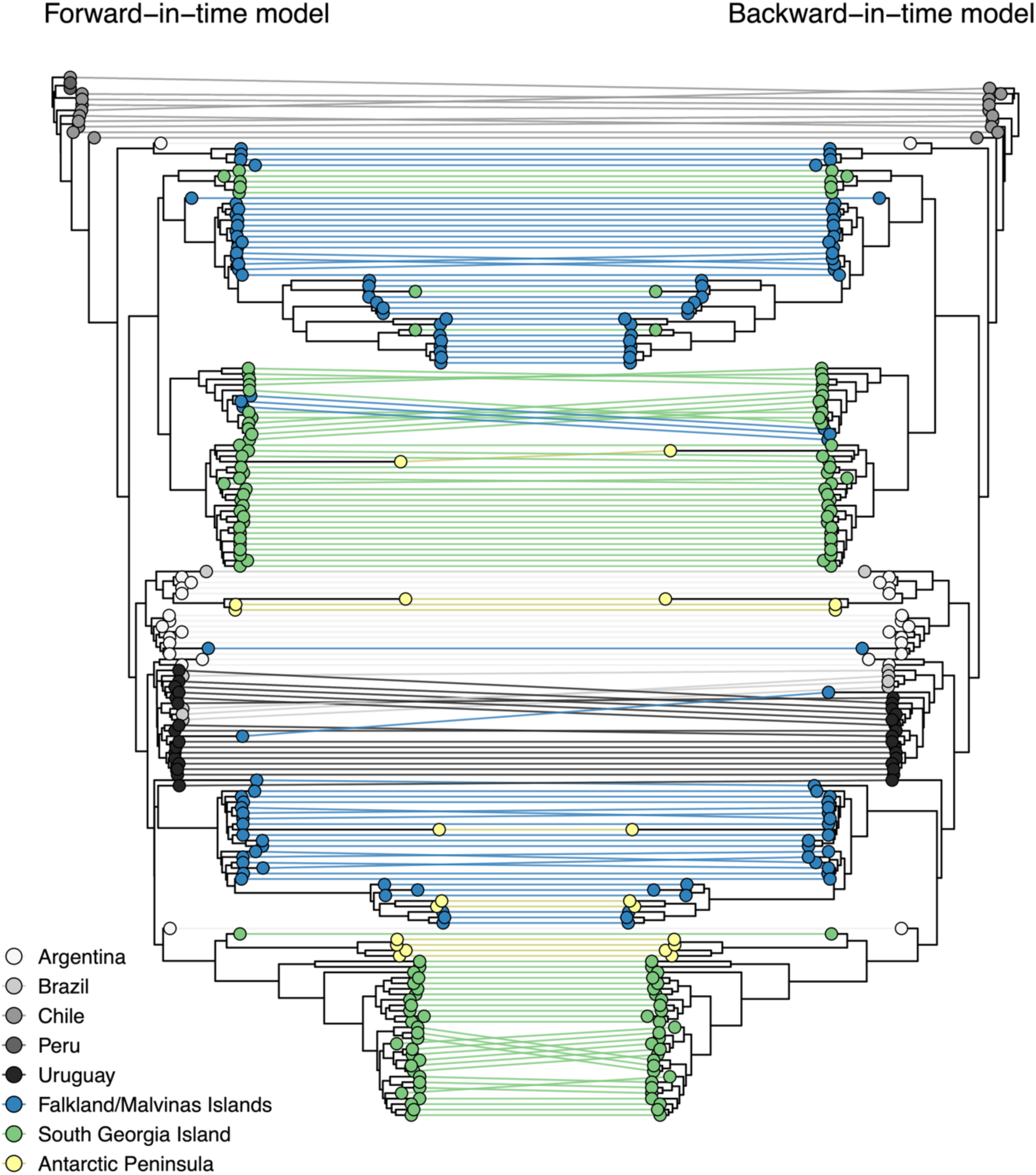
Tanglegram contrasting the majority-rule HIPSTR topologies of Clade II (see Figure 2) under the FIT (left) and BIT (right) phylogeographic models. Minor clustering differences can be observed.

**Supplementary Figure S15.**
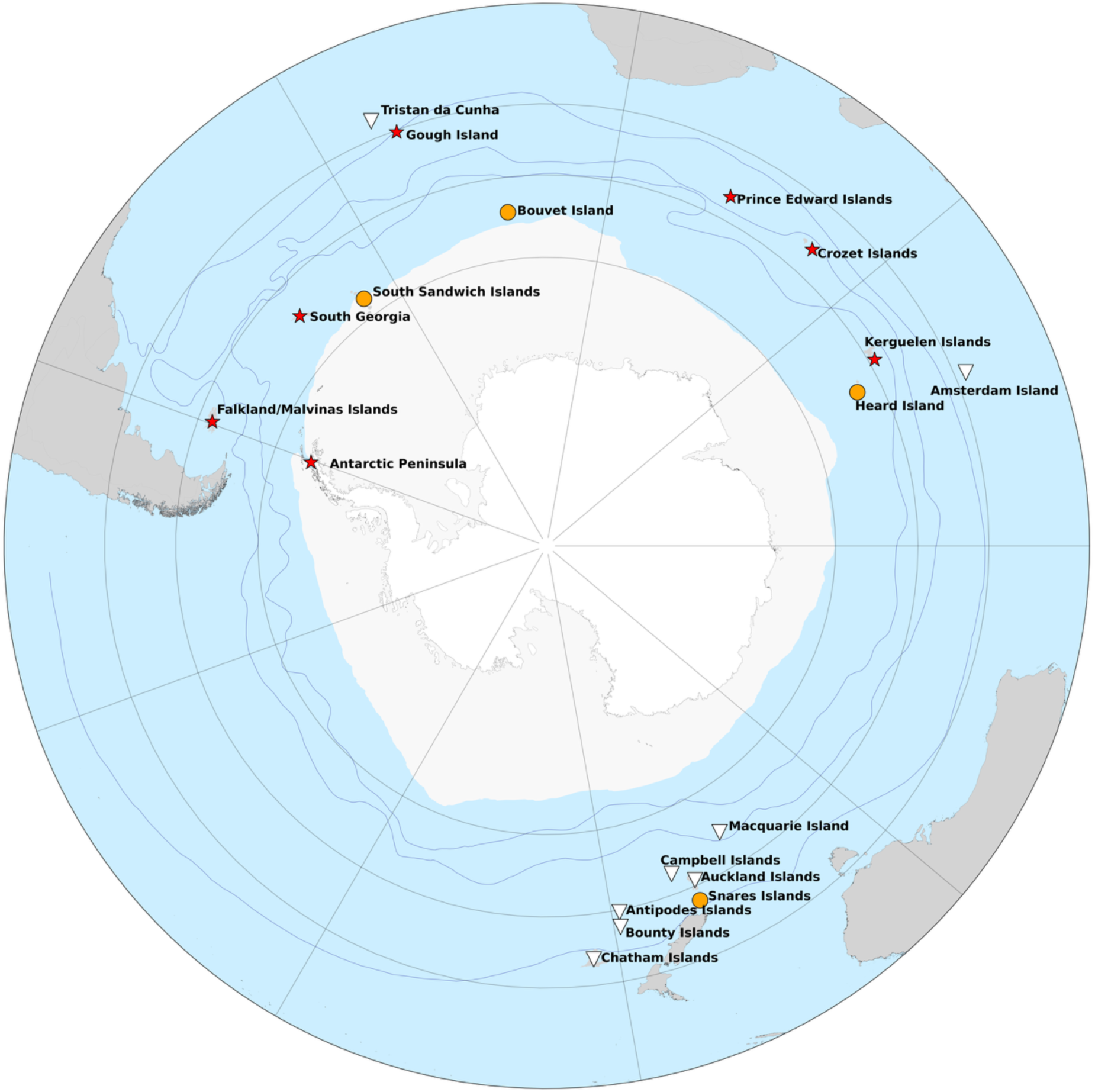
Map of the sub-Antarctic and Antarctica with relevant island names. Islands where HPAI H5N1 has been confirmed are represented by red stars, islands that are thought to be free of HPAI H5N1 are represented by white triangles, and islands that have not been visited since HPAI reached the region are represented by orange circles. On the Antarctic continent and adjacent islands, only the Antarctic peninsula region has reportedly been affected by HPAI H5N1; the rest of the continent remained HPAI-free. The period considered for the HPAI status is the period covered by our analyses, i.e., September 2023 until March 2025, but note that since, HPAIV H5N1 has been confirmed on Heard Island in November 2025 ^99^.

**Supplementary Figure S16.**
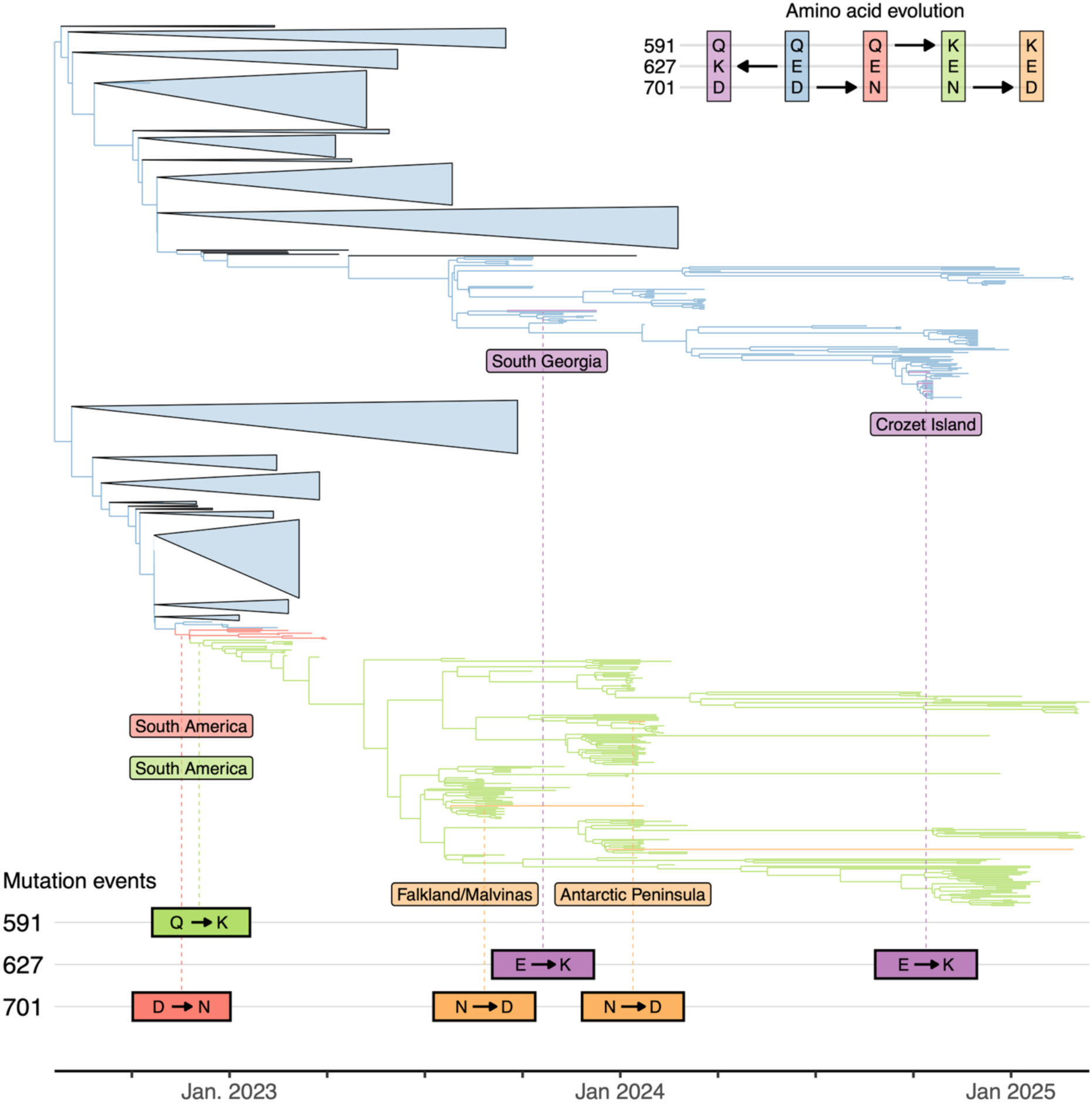
Timeline of inferred mutation events of amino acids 591, 627, and 701 on the full 628-sequence subtree that contains Clades I and II. Sequences outside Clades I and II have been collapsed. Tree branches are coloured according to amino acid composition (top right). Specific inferred mutation events are shown below the tree.

**Supplementary Figure S17.**
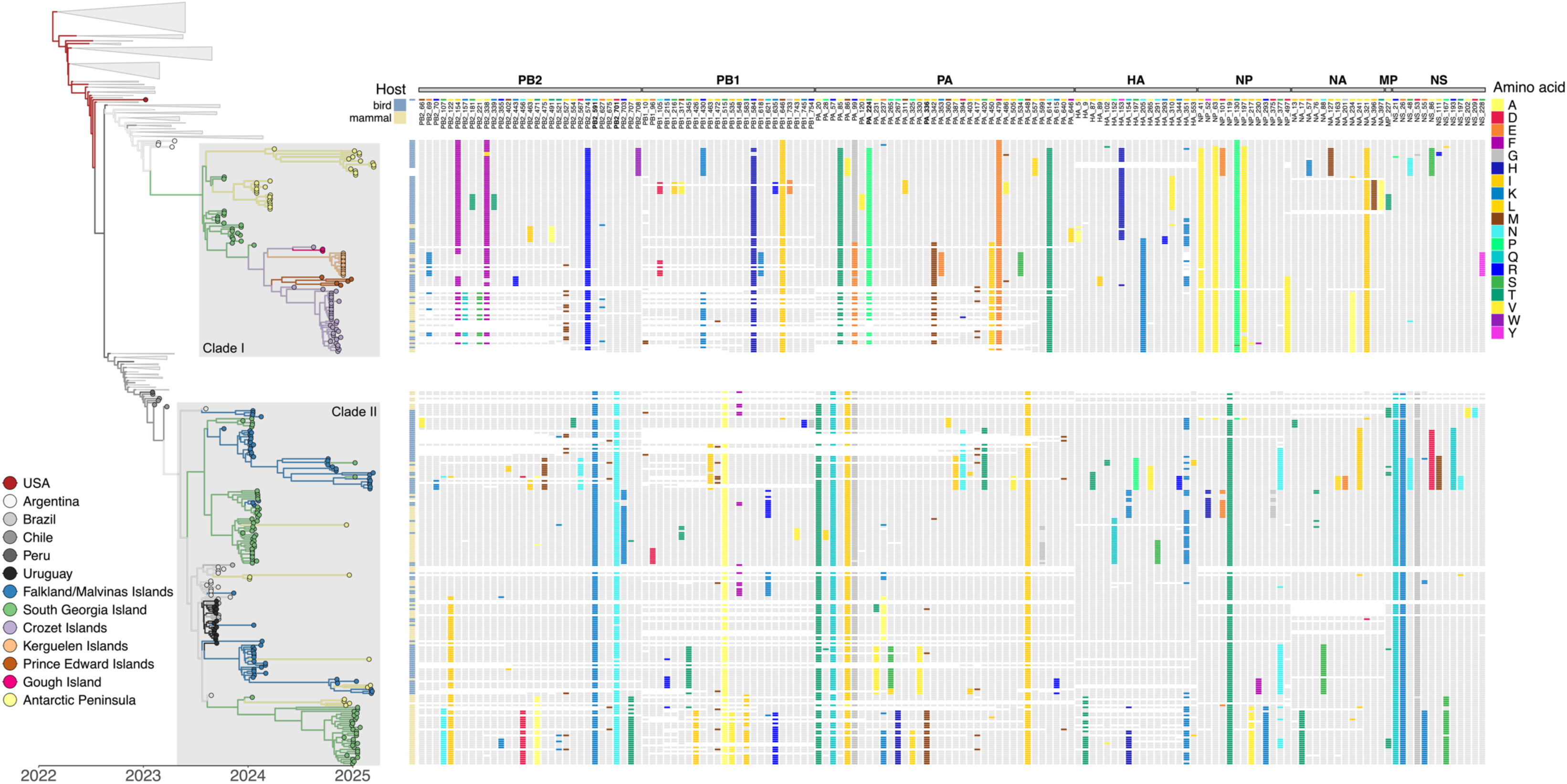
Overview of mutational differences between the predicted sequences in Clade I and II of the proteins PB2, PB1, PA, HA1, HA2, NP, NA, M1 and NS1 coded by the HPAIV H5N1 2.3.4.4b genome. Mutations with at least two occurrences are depicted. The mutation indexing, at the top of the figure, is based on the PB1, PB2, PA, mature H5, NA, NS1, NP and MP.

